# Local optogenetic NMYII activation within the zebrafish neural rod results in long-range, asymmetric force propagation

**DOI:** 10.1101/2024.09.19.613826

**Authors:** Helena A Crellin, Chengxi Zhu, Guillermo Serrano-Nájera, Amelia Race, Kevin O’Holleran, Martin O Lenz, Clare E Buckley

## Abstract

How do cellular forces propagate through tissue to allow large-scale morphogenetic events? To investigate this question, we used an *in vivo* optogenetic approach to reversibly manipulate actomyosin contractility at depth within the developing zebrafish neural rod. Contractility was induced along the lateral cortices of a small patch of developing neural epithelial progenitor cells, resulting in a shortening of these cells along their mediolateral axis. Imaging the immediate response of surrounding tissue uncovered a long-range, tangential, and elastic tissue deformation along the anterior-posterior axis. Unexpectedly, this was highly asymmetric, propagating in either the anterior or the posterior direction in response to local gradients in optogenetic activation. The degree of epithelialisation did not have a significant impact on the extent of force propagation via lateral cortices. We also uncovered a dynamic oscillatory expansion and contraction of the tissue along the anterior-posterior axis, with wavelength within the range of rhombomere length. Together, this study shows dynamic and wave-like propagation of force along the anterior-posterior axis. It also suggests that cell generated forces are actively propagated over long distances within the tissue, and that local anisotropies in tissue organisation and contractility are sufficient to drive directional force propagation.

## Introduction

Long-range force propagation is necessary for morphogenetic processes such as convergence extension, tissue folding and tissue elongation. One important source of subcellular force generation within developing epithelial tissue is the actomyosin complex. The coordinated balance between force generation by this complex and the transmission of this force via cadherin-based cell-cell adhesions is integral to initiating morphogenesis at the tissue scale (Clarke & Martin, 2021; Heisenberg & Bellaiche, 2013). Specific patterns of subcellular force at the apical surfaces and apical-lateral junctions have been shown to facilitate morphogenesis at the whole embryo scale. For example, the anisotropic contractility of planar-polarised actomyosin enables *Drosophila* germband extension (Bertet *et al*, 2004; Dicko *et al*, 2017; Rauzi *et al*, 2010; Simoes Sde *et al*, 2010; Zallen & Wieschaus, 2004). However, transmission of forces between the lateral sides of cells is also potentially instrumental in facilitating complex morphogenesis. For example, light sheet imaging of the columnar *Drosophila* germband epithelium revealed that non-muscle myosin II (NMYII)-dependent contractile oscillations occur at points throughout the apical-basal axis of cells, where they can initiate topology change along the cell length (Vanderleest *et al*, 2024). In pseudostratified epithelia such as the neuroepithelium, each cell is in contact with multiple and differing neighbours along its apical-basal length (Gomez *et al*, 2021). This means that forces transmitted laterally have the potential to transmit to more neighbours than those at the apical surfaces.

Until recently, a major limitation in studying *in vivo* force transmission has been the inability to manipulate mechanics or signalling with sufficient spatiotemporal control. The advent of *in vivo* optogenetic approaches has huge potential to unlock previously unreachable mechanisms in biology. It allows the role of subcellular and cellular events to be investigated at the tissue scale within complex organisms, via non-invasive light patterning (Crellin & Buckley, 2024; Krueger *et al*, 2019). This has started to allow researchers to specifically probe the role of actomyosin during morphogenesis with high spatiotemporal precision. These studies have mainly focused on actomyosin networks within polarised epithelia. For example, optogenetic manipulation of Rho1 activity in the *Drosophila* embryo demonstrated that both apical constriction and basal relaxation are required to enable tissue folding during gastrulation (Izquierdo *et al*, 2018; Krueger *et al*, 2018). It also demonstrated that NMYII is dynamically recruited to junctions under strain (Gustafson *et al*, 2022). A recent study has used similar optogenetic constructs to demonstrate that a balance between contractility and anisotropic subcellular localisation is required for productive cell-cell rearrangement and tissue fluidity during *Drosophila* germ band epithelial elongation (Herrera-Perez *et al*, 2023). Very recently, optogenetic activation of RhoA at lateral membranes has uncovered that apical-basal cell shortening can contribute to tissue furrowing during *Drosophila* gastrulation (Countryman *et al*, 2024).

An exciting advance in our understanding of lateral force propagation between epithelial cells was achieved via optogenetic activation of actomyosin contractility in one cell of a micropatterned doublet, and measurement of the mechanical response of the second (receiver) cell (Ruppel *et al*, 2023). This demonstrated that the receiver cell shows an active contractile response to force from the first cell, which enables signal strength to be maintained over distance. Interestingly, the study also found that the response of the receiver cell was stronger when the force propagation was perpendicular to the axis of mechano-structural polarisation. This active propagation of force was also seen in small cell clusters. However, until recently it has been difficult to directly measure how lateral cell-scale forces propagate at the tissue scale, particularly in deep, physiologically relevant 3D environments. It is also not clear how force propagation might be affected by the cellular state of the tissue – for example, does tangential force propagate differently between cells that are fully epithelial compared with those that are still more mesenchymal in state?

A tractable tissue in which to investigate these questions is the developing zebrafish hindbrain. During zebrafish hindbrain development, neural epithelial progenitor cells elongate and interdigitate along their mediolateral axis. During this process, they undergo a mesenchymal to epithelial transition, forming a polarised midline along the centre of the initially solid organ primordium (Buckley *et al*, 2013), from which two opposing apical-lateral junctional belts emerge (Symonds *et al*, 2020). This precedes the opening of a fluid-filled neural tube via hollowing. Preceding and overlapping with medial-lateral polarisation, the embryo also undergoes convergence and extension, becoming thinner along the mediolateral axis and longer along the tangential anterior-posterior (AP) axis (Williams & Solnica-Krezel, 2020). During this process, the tissue undergoes divergent anterior-posterior expansive strain and medial-lateral compressive strain, coordinated over very large areas (Bhattacharya *et al*, 2021). We aimed to understand how rapid changes in cellular-level forces might underly such larger scale and slower movements at the tissue level. We therefore sought to uncover how forces propagate during neural rod polarisation, at depth within the zebrafish hindbrain. We investigated 1) the extent and direction that cellular force propagates in the tissue, and 2) whether the strain experienced by the tissue in response to cellular force differs over space (anterior-posterior axis) and time (mesenchymal-epithelial state of the cells). To do this, we further developed an *in vivo* optogenetic approach within zebrafish using the Phytochrome system (Buckley, 2019; Buckley *et al*, 2016) to enable reversible control of actomyosin contractility deep within the developing hindbrain. Using a digital mirror device light pattering system coupled with a spinning disc microscope, we induced contractility along the mediolateral aspect of neuroepithelial cells to investigate force propagation via lateral adhesions, independently to the developing apical-lateral junctional belt at the tissue midline.

## Results

### Optogenetic activation of RhoA allows precise and reversible actomyosin activation deep within the neural rod

We first devised an optogenetic approach to activate actomyosin contractility within zebrafish embryos with minimal background activation. We previously demonstrated that the Phytochrome optogenetic system can used within zebrafish embryos, to reversibly control protein localisation with high spatiotemporal resolution (Buckley *et al*., 2016). Here, we generated a stable transgenic zebrafish line, *Tg(ubb:Ath.Pif6-EGFP-arhgef12a)*, (*Tg(Pif6-EGFP-Larg)* for short, driven by a ubiquitous promoter (see methods). This line expresses a fusion protein consisting of the Arabidopsis phytochrome interacting factor Pif6, linked to the zebrafish-specific Dbl homology (DH) domain of the RhoA guanine nucleotide exchange factor (GEF) ARHGEF12a, also known as Leukaemia-associated Rho GEF (Larg). The DH domain of LARG is highly specific for RhoA, B and C and has very high catalytic activity (Jaiswal *et al*, 2011). Optogenetic recruitment of DH-LARG to the cell membrane has previously been used in cell culture to induce ectopic cleavage furrows within non-adherent interphase HeLa cells (Wagner & Glotzer, 2016), which form via plasma membrane flow in response to increased actomyosin contractility (Castillo-Badillo *et al*, 2020).

To achieve high expression levels of a membrane-associated version of Phytochrome B (PHYB, the binding partner of Pif6), we injected mRNA encoding *PhyB-MCh-Caax or PhyB-Caax* into 4-8 cell stage embryos, alongside the chromophore phycocyanobilin (PCB), as previously published (Buckley, 2019; Buckley *et al*., 2016). This approach resulted in robust and reproducible recruitment of Larg to the cell cortex throughout the embryo in response to 640nm light, while Larg was maintained in the cytoplasm in response to 730nm light (Figure 1A & B). Recruitment of Larg to the cell cortex resulted in an increase in RhoA activity (Figure S1A & B) and an increase in NMYII phosphorylation (Figure S1C & D) within zebrafish neural epithelial progenitor cells *in vivo*. Together, this data demonstrates that the Phytochrome optogenetic system can be used to recruit transgenic Pif6-EGFP-Larg to cell membranes within living zebrafish embryos, and that this is sufficient to activate endogenous RhoA signalling and downstream NMYII phosphorylation, essential for actomyosin contractility.

**Figure 1:**
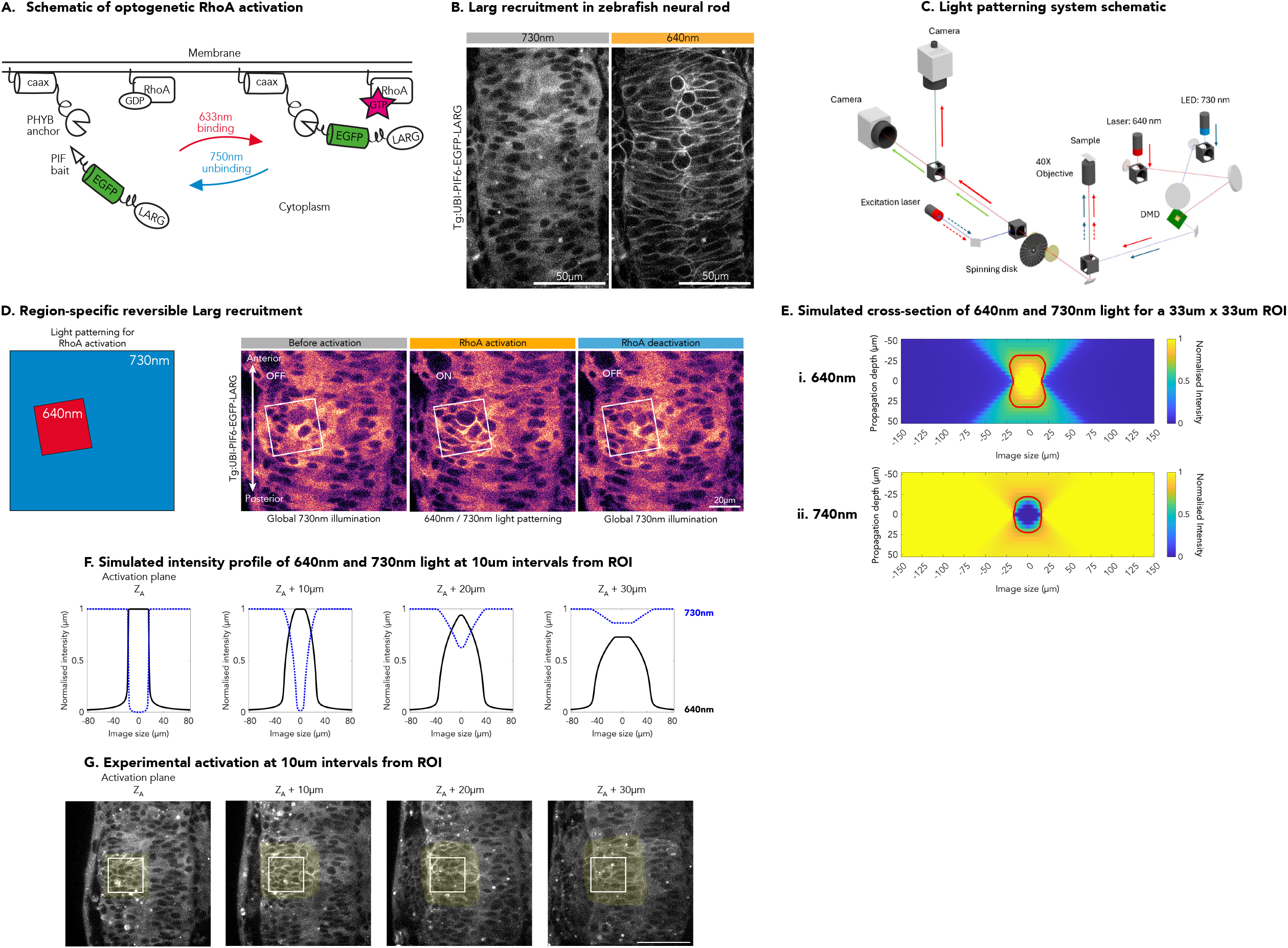
Patterned illumination by DMD results in highly spatiotemporally specific LARG recruitment and derecruitment. A. Cartoon of phytochrome optogenetic system for RhoA activation B. Representative image showing PIF6-EGFP-LARG recruitment to the cell cortex of neuroepithelial cells in 18 somite stage zebrafish neural rod upon 730nm and 640nm illumination. Scale bar 50μm. C. Simplified schematic of the patterned illumination system and the spinning disk confocal microscope. The patterned illumination system is built around a DMD and two light sources (640 nm CW laser and 730 nm LED). D. Diagram of patterned light illumination (640nm in red, 730nm in blue) and representative images showing specificity of recruitment of PIF6-EGFP-LARG to the membrane in the region of activation (ROI, white box). Images were taken at 30µm depth from the surface of the embryo in the neural keel tissue of a 15 somite stage embryo. Scale bar 20µm. E. Simulated cross section (xz) of the 640nm activation (i) and 730nm deactivation (ii) illumination intensity resulting from a 33µm x 33µm ROI. Red line indicates 70% intensity. F. Intensity line profile of simulated activation/deactivation light at activation plane and at 10µm intervals away from the activation plane from a 33µm x 33µm ROI. G. Experimental data demonstrates that activation from a 33µm x 33µm ROI reaches ∼30µm deeper than the imaging plane, with an increasingly broader and weaker region of recruitment (yellow overlay), in line with simulated ∼70% 640nm light. Scale bar 50µm.

We next used a light-patterning approach to restrict Larg recruitment to specific regions of tissue. We built a patterned illumination system, comprising a Cairnfocal digital mirror device (DMD) system coupled to a spinning disc confocal microscope (see methods, Figure 1C). Inverse patterning of activating (640nm) and deactivating (730nm) light resulted in precise and reversible recruitment of Pif6-EGFP-Larg, deep within the neural rod, to specific tissue regions (Figure 1D), with rapid time constants for recruitment and derecruitment (Figure S1E). To predict the shape of this 2D patterned light within the 3D tissue, we used a simplified numerical model based on incoherent superposition imaging modelling (see methods). This predicted that, for an activation region of interest (ROI) of 33µm x 33µm, >70% 640nm recruiting light intensity (∼2.28 mW/cm2) would extend 32µm above and below the imaging plane in an approximately hourglass shape (Figure 1Ei). Lower intensities of 640nm light outside of this ROI would be offset by the inverse patterning of 730nm deactivating light, which has >70% intensity outside of an oval that reaches 22µm (Figure 1Eii). The association of PhyB and PIF6 depends on the ratio between 640nm and 730nm light which is predicted to decrease away from the activation plane (Figure 1F). Supporting these predictions, we found that weaker and broader levels of optogenetic recruitment were present 30µm below the activation plane during *in vivo* experiments (Figure 1G). Similar shapes of illumination were predicted for smaller ROIs (15µm x 15µm), but these had a more restricted axial extent of optogenetic activation (14 µm for >70% 640nm recruiting light) (Figure S2).

Together, this data demonstrates that the Phytochrome optogenetic system can be used to reversibly recruit Larg (and therefore to activate RhoA and phosphorylate NMYII) deep within the developing zebrafish brain. It also demonstrates that a 2D light patterning approach via DMDs can activate tissue in 3D with predictable dimensions.

### A local increase in tissue contractility resulted in long-range, asymmetric tissue displacement

We next investigated whether local NMYII phosphorylation, mediated by optogenetic RhoA activation, caused contractility of the actomyosin cytoskeleton. Within neural keel-rod stage embryos, we activated a 33µm x 33µm ROI on one half of the developing neural tissue at 30µm below the surface, such that the majority of the medio-lateral length of cells was incorporated within the ROI. 14 ± 2.65 cells were contained within the ROI at a single plane (mean ± SD, n = 9). According to our earlier simulation and experimental verification (Figure 1E-G), the total volume of the 640nm illumination would activate ∼144 cells in a ∼60µm z-depth. Locally activating RhoA (and therefore NMYII phosphorylation) within this ROI for 2 minutes caused reversible changes in tissue shape (Figure 2A and Movie 1), and an inward deformation of the basal edge of the tissue in the majority of embryos (Figure 2B-C). This suggests that cells in the ROI are undergoing shortening along their long, mediolateral axis in response to NMYII phosphorylation (Figure 2D). To maintain cell volume, our prediction was that there would be a compensatory tangential anterior-posterior expansion of the region, displacing tissue in both directions of the AP axis (Figure 2E).

**Figure 2:**
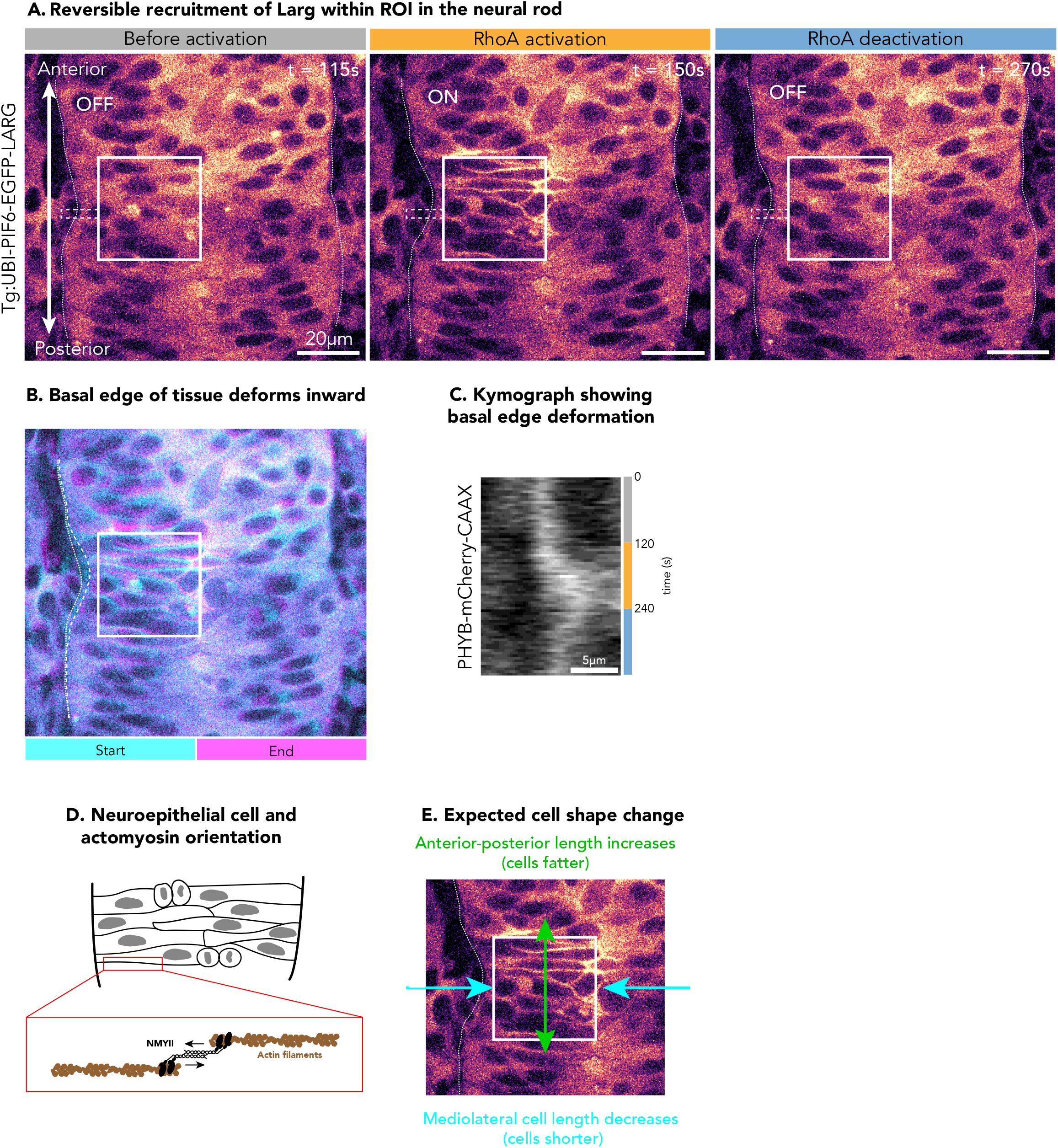
RhoA activation induces local tissue deformation. A. Representative images showing reversible deformation of the basal edge of the tissue following RhoA activation in an 18 somite stage embryo. Solid white box indicates ROI. Dotted line indicates basal edge of the tissue. Dashed box indicates region taken for kymograph in C. Scale bar 20μm. See movie 1. B. Temporal-colour coded image showing changes in cell shape from 100s of activation. Start (cyan) is t = 20s after activation and end (magenta) is t = 120s after activation. Dotted line indicates basal edge of tissue at start and dashed line indicates basal edge of tissue at end. C. Kymograph of the dashed region in A showing the deformation of the basal edge of the tissue. D. Diagram illustrating orientation of neuroepithelial cells and mediolateral shortening of actomyosin cytoskeleton. E. Diagram showing expected cell shape changes during actomyosin activation within the ROI.

To test this, we measured the effects of this local increase in tissue contractility on the wider surrounding tissue, using particle image velocimetry (PIV) analysis to extract velocity vectors (see methods). Unexpectedly, this uncovered an asymmetric tissue displacement along the anterior-posterior axis in a wide area of surrounding tissue. During RhoA activation, displacement occurred tangentially to the long axis of the activated cell and either in the anterior or in the posterior direction. Tissue movement reversed during RhoA deactivation, demonstrating that there was a degree of elasticity within the system (Figure 3A, Movies 2-3). Asymmetric propagation of velocity was highly reproducible, as seen in kymographs depicting mean embryo velocity over time and space (Figure 3B). Increased velocity propagated at least 84 µm towards the anterior or 56 µm towards the posterior during peak velocity (Figure 3C).

**Figure 3:**
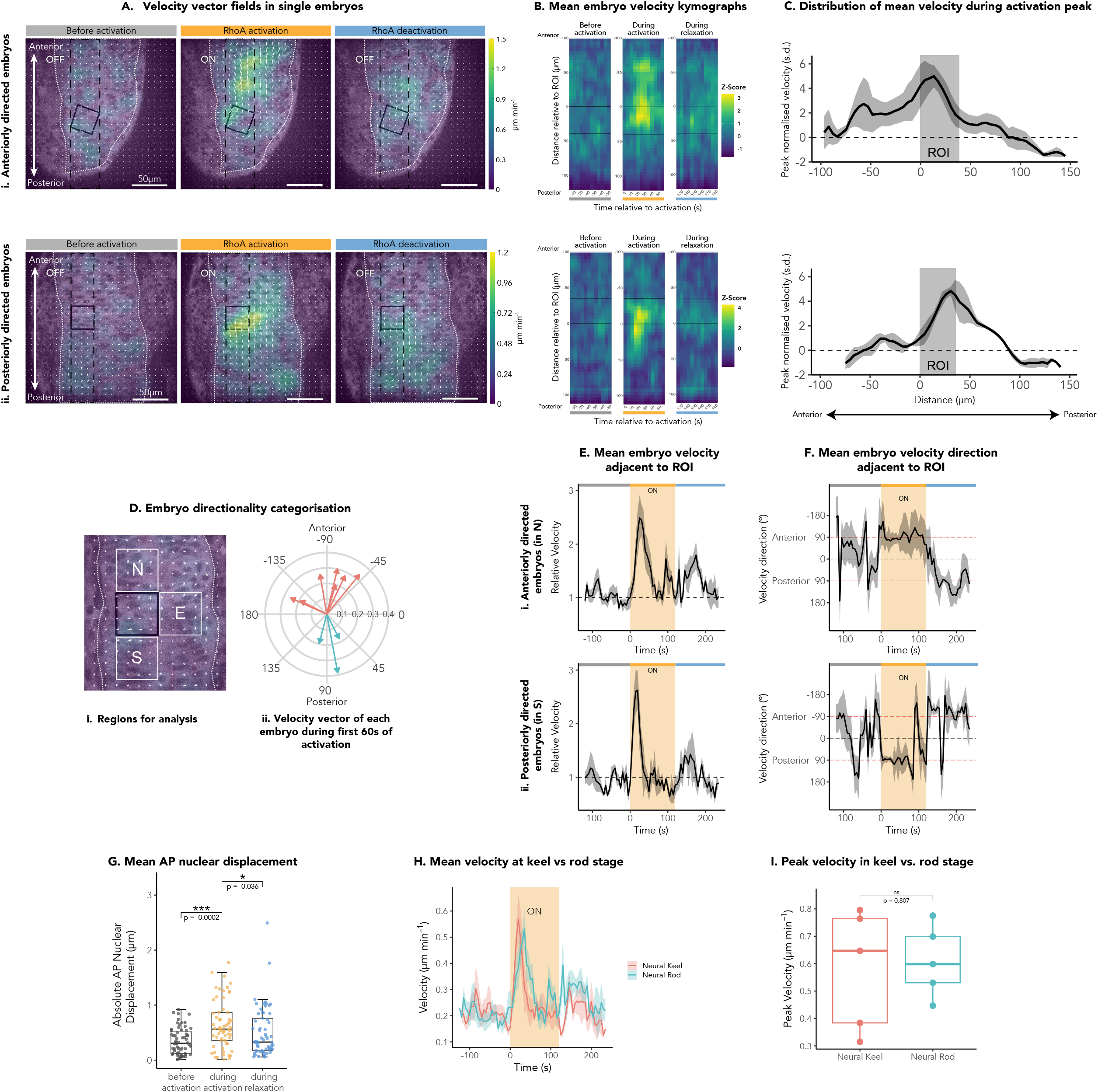
Local RhoA activation induces a long-range, asymmetric tissue displacement. A. Velocity vector fields of neural keel stage embryos before activation, during peak activation and after activation. i. Anteriorly directed (see Movie 2). Ii. Posteriorly directed (see Movie 3). B. Space-time heatmaps of normalised velocity. Each panel represents an anterior-posterior column (see black dashed line of A) overlying the area of activation (ROI, solid black lines) and covers a 55s period, either before activation, immediately upon RhoA activation and immediately post activation. Normalised velocity is in z-scores / standard deviations. i. Average of anteriorly directed embryos, n = 7. ii. Average of posteriorly directed embryos, n = 3. C. Average normalised velocity in the anterior-posterior column overlying the ROI at the time of peak velocity for anteriorly directed embryos (i, n = 6) and posteriorly directed embryos (ii, n = 3), showing that the increase in velocity propagates along the anterior-posterior axis in the anterior (i) or posterior (ii) direction. Normalised velocity is in z-scores / standard deviations. D. Categorisation of embryo directionality. i. Diagram showing the regions analysed adjacent to the ROI. N represents ‘north’, a region anterior to the ROI, S represents ‘south’, a region posterior to the ROI, and E represents ‘east’, a region medial to the ROI. ii. Polar plot showing the weighted average velocity vector within the combined NSE regions during the first 60s of RhoA activation. Each vector is one embryo and colour indicates anteriorly (red) or posteriorly (blue) directed embryos. The radius shows velocity magnitude (µm min^-1^). E. Average normalised velocity (relative to the preactivation mean). i. in the ‘north’ region of anteriorly directed 12-21 somite stage embryos, n = 6. ii. in the ‘south’ region of posteriorly directed 13-15 somite stage embryos, n = 3. F. Average velocity direction (relative to the preactivation mean). i. in the ‘north’ region of anteriorly directed 12-21 somite stage embryos, n = 6. Demonstrates a switch in mean direction towards the anterior during activation, and towards the posterior during deactivation of RhoA. ii. in the ‘south’ region of posteriorly directed 13-15 somite stage embryos, n = 3. Demonstrate a switch in mean direction towards the posterior during activation, and towards the anterior during deactivation of RhoA. G. Absolute nuclei displacement in the AP axis in the ‘north’ or ‘south’ region of anteriorly or posteriorly directed embryos respectively, n = 62 nuclei from 10 embryos from 6 experiments. Anterior-posterior nuclear displacement of 0.639 ± 0.435 µm was seen after 60 seconds of RhoA activation, and 0.495 ± 0.448 µm 60 seconds post activation (mean ± SD). A statistical significant difference between groups was found using the Friedman test (X2(2) = 11.6, p = 0.00296) with significant differences between before and during (***p = 0.0002) and during and after (*p = 0.036) RhoA activation using the paired Wilcoxon signed-rank test with Bonferonni correction. Boxplot shows median, points represent nuclei. H. Average velocity in the ‘north’ or ‘south’ region of neural keel (12-15 somite stage embryos, n = 6) and neural rod (16-21 somite stage embryos, n = 4). I. Peak velocity of in the ‘north’ or ‘south’ region of neural keel (12-15 somite stage embryos, n = 6) and neural rod (16-21 somite stage embryos, n = 4) is not significantly different. The peak velocities were not significantly difference using the two sample t test (t(8) = 0.111, p = 0. 915). Boxplot shows median, points represent embryos.

For each responding embryo (see methods), the mean vector direction was calculated from tissue immediately anterior (north), posterior (south) and medial (east) to the region of RhoA activation (Figure 3D). This was used to categorise the response as anteriorly or posteriorly directed. We then analysed mean velocity magnitude and direction in the tissue immediately north or south to the area of RhoA activation, categorised by direction as above. This demonstrated a rapid and highly reproducible response, regardless of direction, with a mean peak in relative velocity of 2.56 times baseline following 22.5s of RhoA activation. This was followed by recoil of the tissue upon RhoA deactivation, characterised by a second peak in relative velocity of 1.61 times baseline at 48s post-deactivation (Figure 3E). Similarly, vector direction change was also highly reproducible, clearly switching between anterior or posterior directions at the point of RhoA activation/deactivation, demonstrating the tissue moving away from the ROI during activation and recoiling back during deactivation (Figure 3F). Conversely, little change in velocity magnitude and a lack of coordinated direction was seen on the opposite side of the ROI (south for anteriorly directed embryos or north for posteriorly directed embryos) (Figure S3).

The rapid timescale of this response indicates an immediate displacement of surrounding tissue in response to a change in contractility, not a neighbour exchange, which would likely occur on a longer timescale (minutes, rather than seconds) (Das *et al*, 2021; Dawney *et al*, 2023). In support of this hypothesis, we found that RhoA activation induced a significant and reversible anterior-posterior nuclear displacement of approximately 0.5 µm (Figure 3G). Therefore, a small-scale nuclear displacement of ∼0.5 µm was propagated a large-scale tissue distance along the AP axis (at least 56-84 µm). Interestingly, the velocity magnitude was not significantly different between neural keel (12ss-15ss, still undergoing the mesenchymal-to-epithelial transition) and neural rod (16ss-21ss, fully epithelial) stage embryos (Figure 3H-I). This suggests that force propagation between lateral cell-cortices was not affected by the epithelial state of the tissue. We therefore continued to pool neural keel and rod embryos for further analysis. Together, this data demonstrates first that locally activating RhoA optogenetically is an effective method to cause local tissue contractility at depth within a 3D tissue. Second, it uncovers an unexpectedly asymmetric and long-range displacement of surrounding tissue along the AP axis, that is independent of the epithelial state of the tissue.

### Propagating spatial oscillations of expansion and contraction are present along the AP axis

To further investigate the tissue shape-change during this asymmetric displacement of the tissue, we extracted strain rate tensors from the PIV data for anteriorly directed embryos (see methods). We first considered isotropic strain rate, which quantifies the rate of expansion and contraction of the tissue (Figure 4A). Kymographs were generated of isotropic strain rates across an anterior-posterior column overlying the ROI in anteriorly directed embryos (Figure 4B & C). Before actomyosin activation (i.e. in a control situation), this demonstrated the presence of locally propagating spatial oscillations of expansion and contraction along the anterior-posterior axis (Figure 4C and S4A). Upon actomyosin activation, a pronounced area of expansion was seen within the ROI (Figure 4C). This represents the anterior-posterior expansion of the tissue during activation in response to mediolateral cell shortening (Figure 2E). The spatial oscillation of expansion and contraction along the AP axis was temporally synchronised during activation, creating a static standing wave along the anterior-posterior axis (Figure 4C-D and S4B). During relaxation of actomyosin, the polarity of this response was reversed, with contraction seen within the ROI and expansion either side (Figure 4C) and temporal synchronisation was lost (Figure S4C). The magnitude of the isotropic strain rate was relatively low (Figure 4H), with a mean amplitude of 1.07×10^−4^ ± 2.7×10^−5^ s^-1^ (mean ± SD) across the whole experimental time course, which did not significantly change during activation (Figure 4I). The AP length of the ROI was 30.8-44µum (depending on its angle in the tissue). This was significantly smaller than the mean wavelength of the oscillation (52.8 ± 2.34 µm, mean ± SD), which also did not significantly change during activation (Figure 4J). The ROI is therefore unlikely to have affected the wavelength of oscillation. Interestingly, anterior-posterior rhombomere length ranges between about 50-70 µm (Figure S4D and Maves *et al*, 2002; Terriente *et al*, 2012), suggesting that segmentation of the hindbrain may control the wavelength of oscillation. In posteriorly directed embryos, we found similar oscillations in isotropic strain rate to anteriorly directed embryos, including expansion in the ROI during peak activation velocity (Figure S5A & C). In this case, tissue was mostly expanding anterior to the ROI and mostly contracting posterior to the ROI. This could represent a difference in response between embryos with different directions of tissue displacement. However, the low number of embryos in which tissue moved posteriorly makes it difficult to deduce whether this difference is representative.

**Figure 4:**
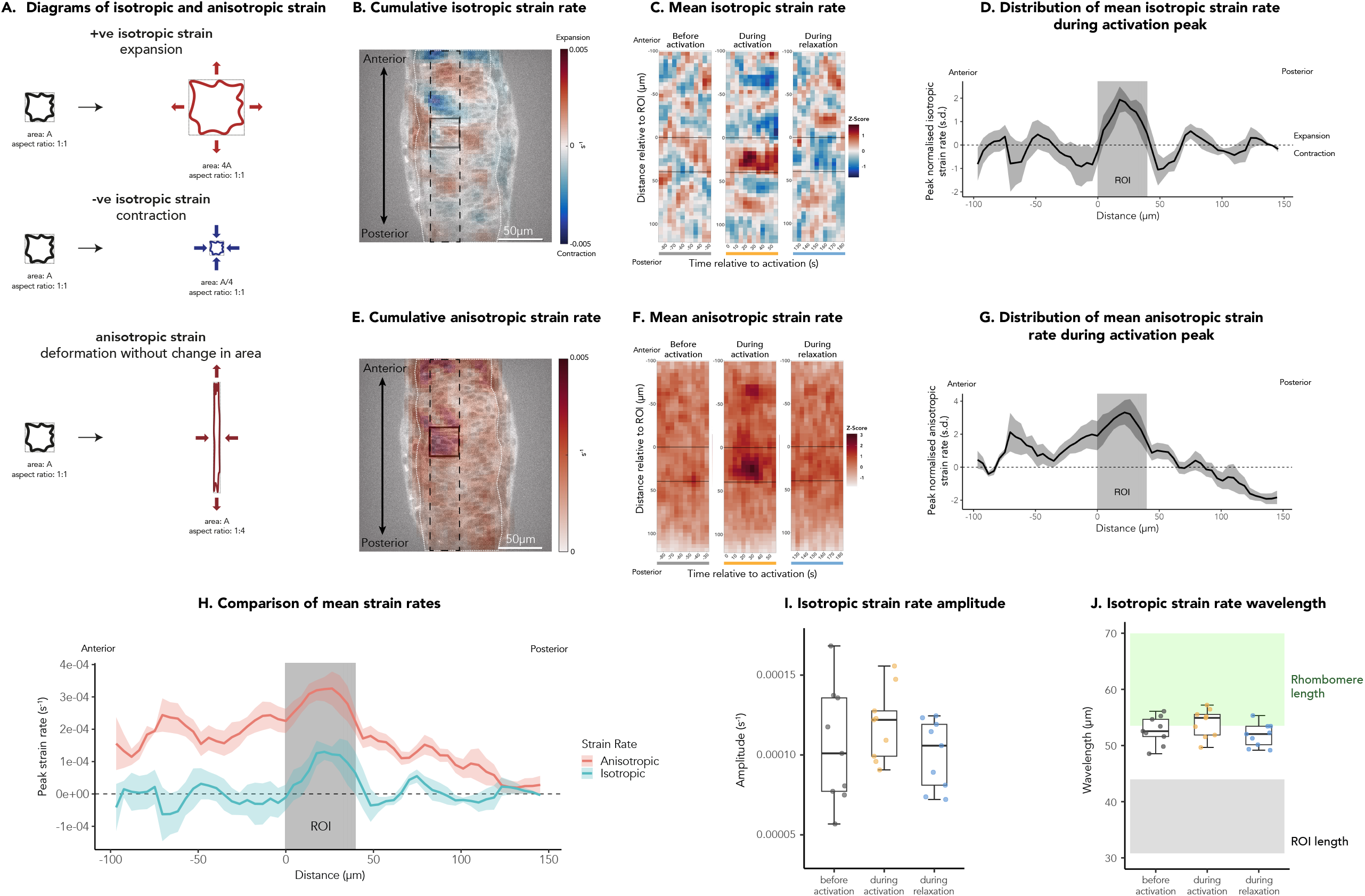
Oscillatory tissue expansion and contraction is sustained during RhoA activation and shear deformation is asymmetrically propagated. A. Isotropic strain rate quantifies the rate of expansion. This is the rate of deformation that leads to a change in area. Positive isotropic strain rate indicates an expansion and negative isotropic strain rate indicates a contraction. Anisotropic strain rate quantifies the rate of shear strain. This is the rate of deformation that leads to a change in shape without a change in total area, where the axes of contraction and extension are orthogonal to each other but can be in any direction (see methods). B. Cumulative isotropic strain rate during the first 55 seconds of activation. Representative image of an anteriorly directed 18 somite stage embryo. C. Space-time heatmaps of normalised isotropic strain rate (average of anteriorly directed embryos, n = 7). Each panel represents an anterior-posterior column (see black dashed line of A) overlying the region of activation (ROI, solid black lines) and covers a 55s period, either before activation, immediately upon activation and immediately post activation. Normalised strain rate is in z-scores / standard deviations. D. Average normalised isotropic strain rate in the anterior-posterior column at the time of peak velocity during activation (average of anteriorly directed 12-21 somite stage embryos, n = 6). Normalised strain rate is in z-scores / standard deviations. E. Cumulative anisotropic strain rate during the first 55 seconds of activation. Representative image of an anteriorly directed 18 somite stage embryo. F. Space-time heatmaps of normalised anisotropic strain rate (average of anteriorly directed 12-21 somite stage embryos, n = 7). Each panel represents an anterior-posterior column (see black dashed line of C) overlying the region of activation (ROI, solid black lines) and covers a 55s period, either before activation, immediately upon activation and immediately post activation. Normalised strain rate is in z-scores / standard deviations. G. Average normalised anisotropic strain rate in the anterior-posterior column at the time of peak velocity during activation (average of anteriorly directed 12-21 somite stage embryos, n = 6). Normalised strain rate is in z-scores / standard deviations. H. Raw anisotropic (pink) and isotropic (blue) strain rates in the anterior-posterior column at the time of peak velocity during activation showing that anisotropic strain has greater magnitude than isotropic strain (average of anteriorly directed 12-21 somite stage embryos, n = 6). I. Mean isotropic strain rate amplitude in 60s of before, during and after RhoA activation. Boxplot shows median, points represent embryos (n = 9, both anteriorly and posteriorly directed embryos). One-way repeated measures ANOVA test showed no significance effect of RhoA activation on amplitude, (F(2, 18) = 2.62, p = 0.1). J. Mean isotropic strain rate wavelength in 60s of before, during and after RhoA activation. Boxplot shows median, points represent embryos (n = 9, both anteriorly and posteriorly directed embryos). The green panel represents the measured range of the anterior-posterior length of rhombomeres 4 and 5 in neural rod embryos (See Figure S5). The grey panel represents the range of anterior-posterior length of the ROI. One-way repeated measures ANOVA test showed no significance effect of RhoA activation on wavelength, (F(2, 18) = 1.87, p = 0.183).

Together, this data suggests that there is an intrinsic oscillatory property to the tissue along the anterior-posterior axis and that rhombomere length may control the wavelength.

### Local actomyosin contractility results in asymmetric shear along the AP axis

Next, we considered anisotropic strain rates which describe the rate of shear (Figure 4A). Kymographs of anisotropic strain rate demonstrated a pronounced region of shear within the ROI during actomyosin activation (Figure 4E-F). Similarly to velocity, the shear strain rate above baseline propagated asymmetrically in the anterior-posterior axis to at least 84 µm in the anterior (Figure 4G) and at least 66 µm in the posterior direction (Figure S5B). We found that during peak velocity, anisotropic strain rate was greater in magnitude than isotropic strain rate, indicating that more anisotropic deformation in shape than changes in area occurred (Figure 4H). Interestingly, a degree of oscillation was also seen in anisotropic strain rate during ROI contraction. Outside the ROI, areas of highest anisotropic strain rate correlated with areas of highest contraction (negative isotropic strain rate) (Figure 4H, S5C).

We next investigated what might cause the asymmetry in tissue response. First, we investigated whether the initial parameters of tissue strain influenced directionality. We did not find a significant difference in strain rate magnitude or in velocity direction before activation between the anteriorly or posteriorly directed embryos (Figure 5A), suggesting that pre-existing strain within the tissue was not responsible for the directionality of response.

**Figure 5:**
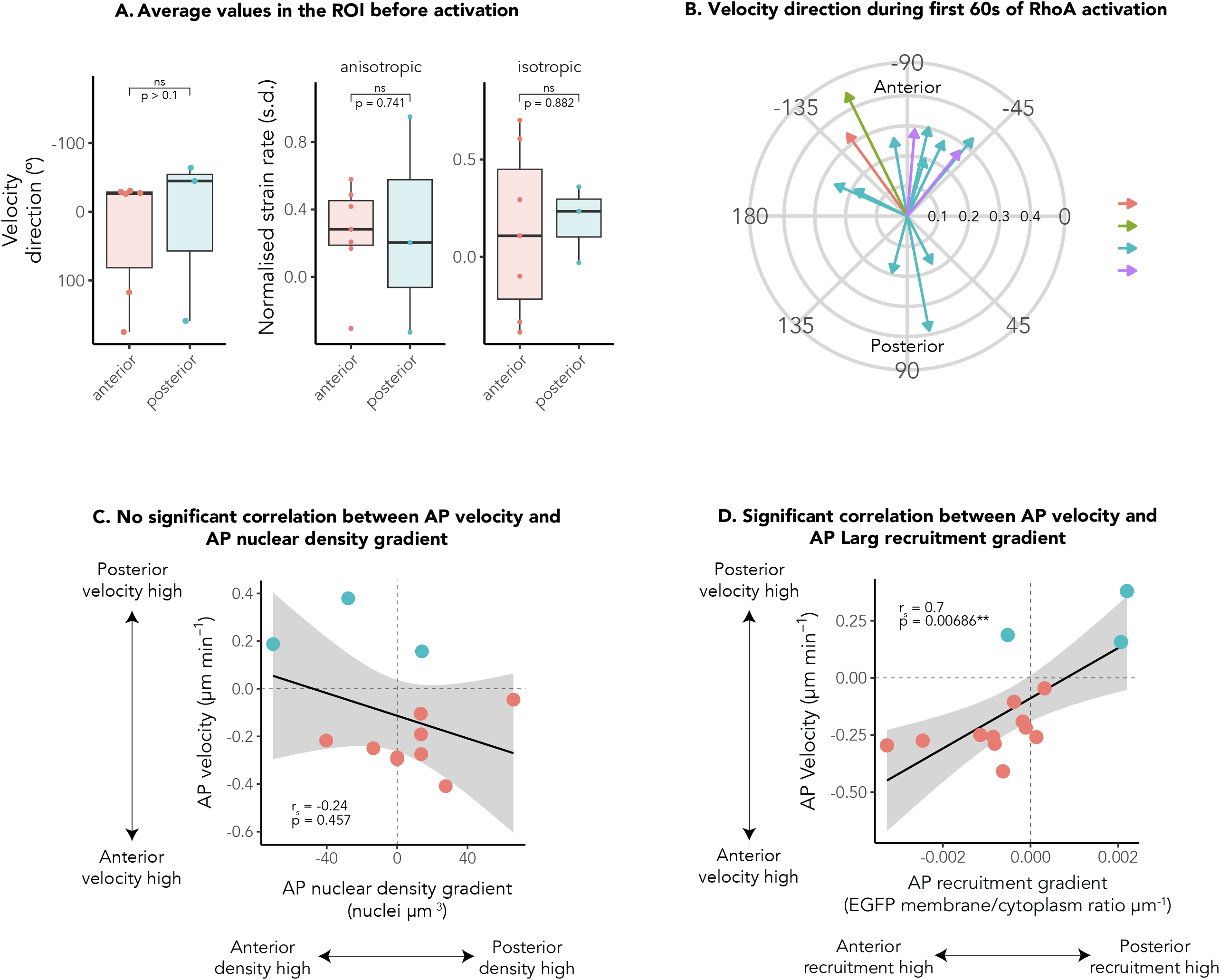
Asymmetric response correlates with gradients in RhoA activation. A. Average velocity direction and average normalised anisotropic and isotropic strain rates in the ROI before activation. Boxplots show median and upper/lower quartiles. Points represent embryos (n = 6 and 3 for embryos that responded towards the anterior and posterior respectively). Watson two sample test (velocity direction, W = 0.1235, p > 0.10) and two-sample t tests (anisotropic and isotropic strain rate, (t(7) = -0.344, p = 0.741 and t(7) = -0.154, p = 0.882 respectively) show no significant difference between the anterior (pink) and posterior (blue) directed embryos. B. Polar plot showing the weighted average velocity vector within the combined NSE regions during the first 60s of activation. Each vector is one embryo and colour indicates rhombomeres. The radius shows velocity magnitude (µm min^-1^). C. Directional anterior-posterior velocity against anterior-posterior nuclear density gradient (compared between tissue neighbouring the north and south sides of the ROI). Positive velocity indicates a posterior direction, and negative velocity indicates an anterior direction. Positive nuclear density gradient indicates higher nuclear density in tissue south of the ROI, whilst negative nuclear density indicates higher nuclear density in the tissue north of the ROI. Points represent embryos that responded towards the anterior (pink) and posterior (blue). Black line indicates linear regression and ribbon indicates 95% confidence interval. Spearman’s rank correlation coefficient showed a weak negative relationship between anterior-posterior velocity and nuclear density, but this was not statistically significant (r_s_ (11) = -0.24, p = 0.457). D. Directional anterior-posterior velocity against anterior-posterior recruitment gradient of LARG within the ROI. Positive velocity indicates a posterior direction, and negative velocity indicates an anterior direction. Positive recruitment gradient indicates higher recruitment of LARG towards the posterior side of the ROI and negative recruitment gradient indicates higher recruitment of LARG towards the anterior side of the ROI. Points represent embryos that responded towards the anterior (pink) and posterior (blue). Black line indicates linear regression and ribbon indicates 95% confidence interval. Spearman’s rank correlation coefficient showed a significant positive correlation between anterior-posterior velocity and recruitment gradient of LARG (r_s_ (11) = 0.7, p = 0.00686**).

Previous work has shown that, during convergent extension, anterior neuroepithelial tissue migrates towards the anterior, while posterior tissue migrates towards the posterior (Bhattacharya *et al*., 2021; Keller *et al*, 2008; Williams & Solnica-Krezel, 2020). To determine whether there was underlying bias in the tissue along the anterior-posterior axis, we next investigated whether the positioning of the region of actomyosin activation within different rhombomeres of the hindbrain influenced the directionality of tissue response. However, the position of the ROI did not influence directionality within the anterior-posterior extent of the hindbrain. For example, tissue in which ROIs were placed in posterior rhombomeres still responded towards the anterior (Figure 5B).

Lastly, we investigated whether anisotropies in tissue structure or RhoA activation in and around the region of interest influenced directionality of the tissue response. There was a non-statistically significant negative relationship between the anterior-posterior nuclear density gradient and anterior-posterior tissue velocity (Figure 5C). We next measured anterior-posterior gradients in LARG recruitment (and therefore RhoA activation) within the ROI (Figure S6A-B). These gradients were significantly positively correlated with anterior-posterior tissue velocity (Figure 5D); LARG recruitment was higher at the anterior side of ROIs within tissue that moved anteriorly, whilst LARG recruitment was higher at the posterior side of ROIs within tissue that moved posteriorly. This analysis also demonstrated that there was a correlation between velocity magnitude and the steepness of the LARG AP recruitment gradient. There was no correlation between the gradient of LARG recruitment and the gradient of the anchor, PhyB-mCherry-Caax, before the timelapse experiment (Figure S6A & C). We therefore suggest that the anterior-posterior gradients in LARG recruitment that we see are due to tissue-intrinsic anisotropies in the localisation of LARG within the tissue. For example, we have observed that, under global activating light, LARG recruitment is higher in cells undergoing mitosis and towards the middle of the neural keel and rod (Figure S6D). Similarly, levels of RhoA activation in wild type and LARG+/PCB-embryos were also higher at the midline (Figure S6E). Junctional proteins start to assemble in medial-lateral regions of the tissue from neural keel stages, initially reaching quite far along the lateral edge of cells (Symonds *et al*., 2020). Therefore, despite the lateral placement of ROIs, this could introduce local gradients in LARG recruitment (and therefore actomyosin contractility) within the ROIs, initiating a highly asymmetric tissue response.

Together, this data suggests that there is an underlying wave-like propagation of force between rhombomeres along the anterior-posterior axis, with alternating areas of expansion and contraction. When a local contractile force is introduced on the mediolateral axis, this tangentially propagates shear strain along the AP axis. Local gradients in actomyosin contractility within the ROI initiate a highly asymmetric response, towards either the anterior or the posterior of the embryo.

## Discussion

### Optogenetic method to locally alter tissue stress deep within a living vertebrate embryo

We have used the phytochrome optogenetics system to manipulate actomyosin contractility within specific regions of tissue, deep within the developing zebrafish neural tube (Figures 1 & 2). The low activation intensities and long wavelengths of light required for activation and deactivation of the system resulted in no noticeable bleaching or phototoxicity over the periods of activation used in this study. Although we did not have need of a second visualisation channel for these experiments, the background deactivation 740nm light made it possible to image simultaneously in both green and red, while activating a specific ROI (Movie S1), expanding the potential for using this technique to study smaller scale processes such as neighbour exchange. Whilst the longer wavelengths and high sensitivity of the phytochrome system precluded the use of high energy variable wavelength lasers for two photon activation over predefined 3D volumes, we were able to predict the shape of 3D activation generated from 2D light patterning (Figure 1E-G, S2). The fast dynamics and reversibility of the phytochrome system allowed us to measure the immediate response of surrounding tissue to these local alterations in tissue stress, using PIV analysis (Figures 3 and 4). Both the optogenetic manipulation of local tissue stress and the measurement of strain rates via PIV are non-invasive. The optogenetic recruitment of LARG allows the activation of endogenous RhoA within the working range of the cells. This approach allows the *in vivo*, physiological measurement of force propagation in response to local alterations in tissue stress.

Our optogenetic approach complements other *in vivo* approaches to manipulate and measure stress and strain. For example, elegant studies using the injection of magnetically responsive oil droplets into zebrafish embryos allowed the precise measurement of anisotropic tissue stress over small distances in response to a specified change in stress, demonstrating a fluid-to-solid jamming transition at the tailbud (Mongera *et al*, 2018). This highlighted the importance of local changes in tissue stress in mediating differences in tissue fluidity and directional morphogenesis.

### Long-range force propagation

While viscous neighbour exchange occurs in response to deformations at the minutes-hours timescale, the zebrafish presomitic mesoderm responds to short-scale deformations of seconds to minutes as an elastic tissue (Mongera *et al*, 2023). In line with this finding, our study in the zebrafish neuroepithelium shows an elastic reversal in tissue displacement direction following activation and deactivation of RhoA (Figure 3), by providing a measurement of the displacement of zebrafish neuroepithelial progenitor cells within seconds of a local alteration in stress. Shear strain was demonstrated to propagate over 80µm along the anterior-posterior axis (Figure 4G). Our findings directly demonstrate that local alterations in cellular strain can be rapidly propagated via lateral cell membranes across large distances, enabling coordinated morphogenesis at the tissue scale. This is relevant to understanding the role of force propagation during directional morphogenesis, such as tissue elongation via compaction (Thomson *et al*, 2021). It is also relevant to understand how forces from external tissues can impact morphogenesis. For example, opposing tissue flows in the developing anterior neural plate, established by differential connectivity to the underlying involuting mesendoderm tissue, lead to tissue folding and shaping (Inman *et al*, 2023).

An important area of future research is to understand how mechanical force is decoded by cells to reciprocally alter biochemical signalling (Kindberg *et al*, 2020). In the current study, the elastic and long-range propagation of force within the tissue suggests an active mechanical response of cells to local deformation via their cell-cell adhesions, as predicted by micropatterning optogenetic experiments (Ruppel *et al*., 2023). Active force propagation has been demonstrated within the *Drosophila* endoderm, in which initial Fog signalling-induced NMYII contractility is propagated as a wave through the tissue without further Fog signalling (Bailles *et al*, 2019). At the cell scale, elegant optogenetic experiments in combination with optical tweezers have demonstrated that the actin cytoskeleton is critical in propagating tensile forces around the cell cortex, and that deformation of the cell membrane alone is not sufficient for force propagation (De Belly *et al*, 2023). Therefore, in the context of multicellular tissues, the cadherins linking actin cytoskeletons between cells likely play a critical role in facilitating active propagation of mechanical forces (Pannekoek *et al*, 2019). We found that velocity magnitude of neural keel (12ss-15ss) and neural rod stage embryos (16ss-21ss) in response to local stress alteration were comparable (Figure 3H-I), enabling us to pool results from keel and rod stage embryos. This suggests that the degree of epithelialisation does not have a significant impact on the ability of cells to propagate force via lateral adhesions. This is likely because the level of lateral cadherin-based adhesions and cellular packing remains similar over the course of mesenchymal to epithelial transition of the zebrafish neuroepithelium. Future studies should investigate how cadherin-based lateral cell-cell adhesions actively transduce and propagate forces. It would be interesting to determine whether force propagation might be altered by local differences in levels of cadherins, such as in the tailbud, which has a N-cadherin dependent gradient in yield stress, linked to the extent of extracellular spaces (Mongera *et al*., 2018).

### Asymmetric directionality of force propagation

In addition to demonstrating rapid, long-range force propagation, our study uncovered an unexpected asymmetry of response; tissue displacement occurred in either the anterior or posterior direction (Figure 3), resulting in asymmetric shear strain (Figure 4E-G). We demonstrate that small local AP gradients in Larg recruitment (and therefore in actomyosin contractility) correlate with the directionality and velocity of tangential force propagation (Figure 5D). This suggests that small local gradients in actomyosin activity initiate a highly asymmetric response across a much wider area of tissue. A computational model of *Drosophila* germband extension only enabled tissue elongation with a graded planar polarised actomyosin (Dicko *et al*., 2017), suggesting that the presence of a gradient (rather than a uniform area of contractility) may be key to the directional response that we see. In the same tissue, mechanical feedback at apical-lateral junctions has also been shown to play a role in the establishment of long-range NMYII gradients and tissue flow patterns (Gustafson *et al*., 2022). If the same process of dynamic NMYII recruitment occurs in lateral membranes, this could explain how the introduction of a local gradient in lateral contractility might initiate an active and directional propagation of force propagation over long distances. We also observe deformation preferentially towards the anterior irrespective of the anterior-posterior positioning of the ROI within the hindbrain (Figure 5B). It is not currently clear why this is the case, although it is possible that the whole AP extent of the hindbrain has an existing bias towards the anterior, as a result of convergence-extension movements (Bhattacharya *et al*., 2021; Keller *et al*., 2008; Williams & Solnica-Krezel, 2020).

As well as gradients in contractility, another parameter that can have a profound effect on the directionality of force propagation is local anisotropies in geometry. For example, during *Drosophila* gastrulation, the anteriorly localised cephalic furrow acts as barrier for tissue deformation, therefore orienting the flow towards the posterior (Dicko *et al*., 2017). Whilst our results suggest that rhombomere boundaries may constrain the spatial periodicity of waves of expansion and contraction, they do not prevent the propagation of anisotropic strain, which reaches over 80µm along the AP axis (Figure 4G). Another potentially relevant geometric feature to consider is curvature, both at the tissue scale and at the scale of individual cells, since curvature is important in modifying the mechanical equilibrium across a surface (Dicko *et al*., 2017). The developing zebrafish neuroepithelium is a curved structure, so it is possible that small differences in the placement of ROIs in relation to overall tissue curvature or local cellular curvature could introduce an imbalance in mechanical equilibrium, leading to asymmetric force propagation. As cell segmentation approaches evolve, it will be important to take into account the 3D shape of cells and tissue to interpret the role of geometry on force propagation. However, so far, the full segmentation of complex pseudostratified epithelial cells has been achieved via more manual approaches (Gomez *et al*., 2021).

Taken together with the literature, our work suggests that a combination of local gradients in contractility and robust mechanical coupling between cells drives a far-reaching and highly directional propagation of force at the tissue level. It also demonstrates that transmission of force via the lateral sides of developing epithelial cells plays an important role in directional force propagation. To fully understand the mechanisms behind directional morphogenesis and precise pattern emergence, future research should address the interrelationship between biochemical and mechanical signalling in this context. For example, during chick hindgut formation, imaging studies demonstrated that an initial FGF8 gradient caused contractility of endoderm cells, which then pulled non-contractile cells into the FGF8 region, causing them to contract (Nerurkar *et al*, 2019).

### Compressional waves along the AP axis

Another unexpected finding was the presence of oscillating regions of isotropic expansion and contraction along the AP axis before activation of RhoA (in baseline conditions). These showed locally propagating spatial oscillations (Figure 4C), providing further evidence for mechanical coupling between cells. Upon activation of RhoA within an ROI, areas of expansion and contraction were synchronised in time, creating a standing compressional wave along the AP axis with a mean wavelength of ∼53µm (Figure 4B-D, J, S4B). Temporally oscillating stresses have previously been measured in the zebrafish hindbrain using deformable polyacrylamide beads, and were suggested to result from synchronised cell divisions (Träber *et al*, 2019). In future, it would be interesting to see how these temporal oscillations intersect with the spatial oscillations that we find at the rhombomere length scale. The emergence of such self-organising oscillations of expansion and contraction are strikingly similar in pattern to that of spatiotemporal oscillating cell movement and isotropic strain seen in confined monolayers of human keratinocytes and enterocytes (Peyret *et al*, 2019). This study revealed that the periodicity of oscillations and velocity correlation length correlated with the size of the geometrical confinement of the tissue and that cadherin-based cell-cell adhesions are critical to generate oscillatory behaviour. Such similarities in oscillatory patterns, despite the different behaviours (cell migration vs. cell deformation) and timescale (30 hours vs. a few minutes), highlight the propensity of epithelial tissue for self-organised oscillations (Balaji *et al*, 2017; Fernandez-Gonzalez & Zallen, 2011; He *et al*, 2010; Martin *et al*, 2009; Sawyer *et al*, 2011; Serra-Picamal *et al*, 2012). It also illustrates again the importance of cadherin-based adhesions in propagating force between cells, therefore enabling pattern emergence at the tissue level.

Surprisingly, we found that, whilst isotropic strain rate waves become static during activation, the wavelength of isotropic strain rate across the anterior-posterior axis is sustained (Figure 4J), appearing to be an inherent property of the tissue. This suggests that the spatial periodicity of oscillations may be controlled by the mechanical environment, such as geometrical confinement (Peyret *et al*., 2019). Geometrical constraints in our tissue could include the basal edges of the neural tube and segments such as the rhombomeres, of which the latter has an approximate 50-70 µm AP length (Figure S4D and Maves *et al*., 2002; Terriente *et al*., 2012). The wavelength in isotropic strain is within this range. Therefore, it is possible that spatial periodicity is controlled by rhombomeres, or that the natural frequency of the tissue instructs rhombomere size. This presents an area of future study. We hypothesise that the spatial oscillations in expansion and contraction that we see along the AP axis may be involved in directionally propagating force between rhombomeres along the AP axis.

## Conclusions

We have demonstrated that optogenetic activation of RhoA signalling leads to phosphorylation of NMYII and rapid contraction of the actomyosin cytoskeleton within zebrafish neuroepithelial progenitor cells, *in vivo*. Inverse light patterning of recruiting and derecruiting light in a 2D plane results in a predictable 3D recruitment ROI deep within the developing neural tube. This technique is adaptable to a wide range of research questions, since it allows the behaviour of cells and tissue to be quantified before, during and after a precise and reversible manipulation in actomyosin contractility, deep within the developing zebrafish embryo. The ubiquitous promoter within the transgenic line drives expression of Larg in all tissues. Particle image velocimetry demonstrated an underlying wave-like propagation of forces along the anterior-posterior axis, with oscillations between tissue expansion and contraction within the rhombomere length scale. Local changes in tissue contraction in the mediolateral axis of neuroepithelial progenitor cells led to rapid tangential shear deformation of the tissue along the anterior posterior axis. This force propagation was long ranging, suggesting the active mechanical propagation of cell generated force. The response did not depend on the level of epithelialisation of the tissue, and propagated asymmetrically along the anterior-posterior axis, in response to local AP gradients in actomyosin contractility. This helps to explain how precise morphogenetic patterns can occur across distance in response to relatively small local anisotropies in cellular contractility. Further investigations into the molecular mechanisms of force transmission via cadherin-based cell-cell adhesions will be important to further understand how such anisotropies might arise. Together, the ability to precisely and reversibly manipulate actomyosin contractility at depth within a physiological *in vivo* vertebrate system has uncovered rapid, long-range and directional mechanical communication between cells.

## Methods

### Experimental design

All procedures were carried out with Home Office approval and were subject to the University of Cambridge and the University of Manchester AWERB Ethical Committee review (PPL PF2F15847 and 4544659). Experiments were designed according to ARRIVE guidelines (https://arriveguidelines.org/arrive-guidelines, see supplementary methods).

### Transgenic line details

*Tg(ubb:Ath.Pif6-EGFP-arhgef12a)* fish express the first 100 amino acids of Arabidopsis PIF6 (Gene ID: 825382), codon optimised for mammalian expression, as previously published (Buckley *et al*., 2016). This is fused with EGFP and the majority of the DH domain from zebrafish Arhgef12a (Gene ID: 562455). The fusion protein is expressed under the ubiquitin B promoter, which drives transcription in all tissues. See supplementary methods.

### Patterned illumination system

The Cairnfocal is a commercial system designed and developed by Cairn research (Martin Thomas 2014, Two mirror optical arrangement for digital micromirror device, patent EP2876479B1), with minor custom modifications (as detailed below). The system is composed of a digital mirror device (DMD), two light sources, two sets of Offner triplets and a plurality of individually adjustable mirrors. The layout of this system allows two light sources to be simultaneously patterned by the DMD. To achieve inverse conformation of activating (640 nm) and deactivating (730 nm) light onto the tissue, a LDI-7 Laser Diode Illuminator (89 North) and a 730 nm LED (Thorlabs M730L5) are positioned on the left and right beam path. The light from each source is collimated and projected to the Offner triplet. Both sets of Offner triplets are identical and focus the incoming light on the DMD at the incident angle of positive 12-degree and negative 12-degree.

The DMD used in the system is ViALUX WUXGA 0.96” Type A. The DMD is positioned at the relayed image plane which is centred on the optical axis of the light path to the microscope. Each micro-mirror on the DMD flips to the “on” or “off” position with respect to the input pattern to reflect the light from either source. The patterned beam is re-collimated, folded and projected to the back port of a Nikon Ti2 inverted microscope to activate/deactivate specific areas of the specimen. A shortpass dichroic filter (T610SPxxr, Chroma) is used in the fluorescent turret of the microscope frame to direct the light to the sample. At the sample plane for activation, 1% 640 nm laser power (3.25 mW/cm^2^) and 100% 730 nm LED power (293 mW/cm^2^) were used for patterning experiments. The DMD was controlled using python scripts called through MetaMorph Journals.

### Optogenetic imaging

Embryos were cultured at 24-28.5°C, to achieve the desired stage of development. Following injection with mRNA and PCB, embryos were kept in the dark and mounted under 730-740nm light using mounted and diascopic light sources (Thorlabs, M730L5, and Zeiss, KL1500 LCD fitted with a 740nm band-pass filter (Envin Scientific), respectively). Embryos were mounted in 1-1.5% agarose diluted in E3 medium. The stage positions of embryos mounted on the light patterning microscope were found under 730 nm LED light, to prevent activation of RhoA by background light. The field of view was positioned at 30µm below the enveloping layer and centred over rhombomere 5 of the hindbrain, using the otic vesicles as landmarks. Single dual colour images following 1 minute of global 630nm and 730nm illumination were taken before experiments to verify successful optogenetic recruitment and derecruitment. Embryos were exposed to 2 minutes of global 730nm illumination prior to experiments to ensure experiments began from a deactivated state. For optogenetic timelapse experiments, images were taken every 5 seconds, using 1 second exposure, concurrent with constant light patterning (see Figure S8A). To estimate the level of activation in the z-axis, a 33×33 µm ROI was activated within the neural tissue 30µm beneath the enveloping layer for 2 minutes. Snapshots were the taken at different axial distances from the ROI plane. To ensure that imaging did not cause background optogenetic activation, each snapshot was interspersed with another 2 minutes of light patterning (see Figure S8B). Experimental photoactivation and imaging pipelines were operated using MetaMorph journals.

### 3D activation simulation (incoherent superposition imaging modelling)

To simulate the intensity distribution of the activation and deactivation light around the imaging plane a model based on diffraction-limited imaging theory was used, implemented in MATLAB based on code from Voelz, 2011. An aberration free incoherent superposition imaging model simulates the intensity distribution after propagating through the objective lens at multiple distances from the objective (Voelz, 2011). The objective was set to a 40x silicone oil immersion objective with an NA of 1.25. The image size was set to 330 × 330 µm (1200 × 1200 pixels) to match experimental conditions. The simulations were run for 640 nm light (activation) and 730 nm light (deactivation). Light reflected from the active pixels of the DMD to the objective was assumed to be perfectly collimated and focussing on the ideal imaging plane. The region of interest for activating was either set to the 15µm x 15µm or 33µm x 33µm on the imaging plane. The overall propagation range was +/-50 µm in steps of 5µm. The maximum intensity of the activation light was normalised to 1 and used to reconstruct the 3D volume.

### Particle image velocimetry

Tissue velocity was quantified through particle image velocimetry (PIV) by cross-correlating successive image pairs.

Before analysis, images were manually screened for high specificity of recruitment in the ROI. Images were converted to 8-bit and horizontally flipped or rotated in Fiji to ensure ROIs were on the left side and the anterior-posterior axis was aligned along the y-axis. Each image underwent preprocessing, specifically noise reduction with a 5×5 pixel wiener filter (wiener2 of the Image Processing Toolbox, MATLAB) and a running average of the previous and following frames using a custom MATLAB script.

For PIV analysis, we used PIVLab v2.56 for MATLAB, implementing an initial 64×64 pixel interrogation window and secondary 32×32 pixel window at 50% overlap (equivalent to 17.6×17.6 µm and 8.8×8.8 µm respectively), which are in the order of the size of a cell. Resulting velocity fields were smoothed with a 5×5 vector spatial filter before quantification and further analysis.

### Strain rates

Strain rates characterize local tissue deformation kinetics. They can be decomposed into isotropic and anisotropic components (Serrano Najera & Weijer, 2020). The strain rates can be derived from the gradient of the velocity *v* obtained from PIV. ∇*v* can be decomposed into a symmetric part (strain rate tensor, ξ) and an anti-symmetric part (spin tensor, ω):

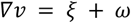

Where:

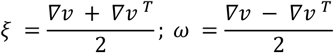

Expanding the strain rate tensor, ξ:

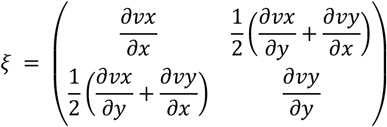

ξ can be further decomposed into an Isotropic Term (ξ_iso_) with the form of a scalar times the Identity, or unit tensor (I), plus an Anisotropic Term (ξ_ani_) with the form of a traceless symmetric tensor (Figure 4A).

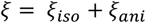

Where:

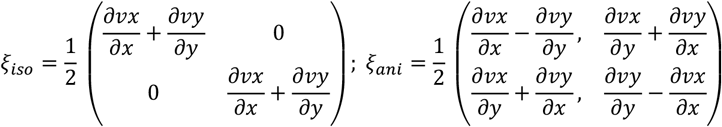

The eigenvalue of isotropic strain rate tensor (ξ_iso_) describes the magnitude of local expansion or contraction rate equally in both axes. The anisotropic strain rate tensor (ξ_ani_) represents the shear strain rate. It possesses two eigenvalues of equal magnitude and opposite sign. The eigenvector paired with the negative eigenvalue defines the axis of contraction. Orthogonally, the positive eigenvalue eigenvector outlines the axis of expansion.

### Analysis of velocity and strain rates

To analyse tissue movements surrounding the ROI for optogenetic activation, images were subdivided into a grid with unit areas matching the ROI illumination area. One grid unit was centred on the ROI. The grid units adjacent to the ROI were analysed and referred to as north (anterior), south (posterior) and east (medial) regions (Figure 3Di). Lateral regions were excluded as the ROI was placed towards the lateral edge of the tissue. For each region, we extracted the average and standard deviation for velocity magnitude, velocity direction and strain rate eigenvalues using a custom MATLAB script. To determine the direction of response, we calculated the average circular mean of velocity direction of the combined north, south and medial regions, with mean direction above the equator being categorised as anterior and below the equator as posterior.

To enable comparison of different embryos, we standardised the parameters extracted from the PIV analysis by calculating z-scores. This normalisation indicates how many standard deviations the value is from the pre-activation mean. To determine whether embryos showed a significant increase in tissue movement, we looked at the peak normalised velocity during RhoA activation in the corresponding region to their categorised direction, for example north for anteriorly-directed embryos and south for posteriorly-directed embryos. A z-score above 2 standard deviations was used as a threshold to filter embryos for further analysis.

Additionally, an anteroposterior column encompassing the width of the ROI was evaluated. Each parameter was averaged across the column width for discrete positions along the embryo anteroposterior axis, using a custom MATLAB script. To analyse the oscillation in isotropic strain rate, data was smoothed with a Savitzky-Golay filter to enable identification of peaks. Peak location was then used to calculate amplitude and wavelength from the raw data.

In summary, quantification focused on local regions immediately surrounding and along the axis of optogenetic activation, providing critical insight into light-induced tissue deformations and movements. Partitioning into discrete grid areas and columnar analysis facilitated quantitative characterization of propagation effects.

### Statistical analysis

Statistical analysis was carried out in R Studio using R version 4.3.2. Standard statistical analyses were carried out using the rstatix package, and circular statistics using the circular package. We performed Shapiro-Wilk tests for normality and Levene’s tests for equality of variance before selecting the appropriate statistical tests. Details of specific hypothesis tests are described in the figure legends.

## Data and resource availability

All data and resources will be made freely available after publication. Custom MATLAB and R scripts are found at https://github.com/Buckley-Lab-opto/Crellin-et-al-2024/. *Tg(ubb:Ath.Pif6-EGFP-arhgef12a)* fish will be available from eZIRC. Plasmids created in this study will be deposited in Addgene. Image data will be deposited in BioImage archive. For the purpose of open access, the authors have applied a Creative Commons Attribution (CC BY) licence to any Author Accepted Manuscript version arising.

## Acknowledgements

Thank you to Sarah Williams for lab management, cloning and genotyping of the *Tg(ubb:Ath.Pif6-EGFP-arhgef12a)* fish. Thank you to Kasia Anton, for her help in cloning the zebrafish specific gene for LARG and to Masa Tada for giving us a plasmid containing the *ubb* backbone. Thank you to the Cambridge Advanced Imaging Centre for help and access to confocal microscopy and to Cairn research for help in building the light patterning system. Thank you to Cambridge University Biomedical Services for looking after zebrafish stocks. Thank you to Michael Casey for helpful discussions on maths and statistics and invaluable support with coding. Thank you to Sarah Woolner, Veronica Biga and Cerys Manning for feedback on the manuscript.

## Author Contributions

**HAC:** Investigation and data curation (optogenetics imaging experiments), methodology (setting up optogenetics pipeline), formal analysis and visualisation (writing R-code, PIV analysis, designing analysis methods and visualisations, making figures), writing – writing, review and editing.

**CZ:** Investigation and methodology (building the light patterning system, writing code to run the experiments), formal analysis & visualisation (Figure 1E&F, S2).

**GSN:** Investigation, formal analysis, visualisation & methodology (writing Matlab code, PIV analysis and designing analysis methods and visualisations).

**AR:** Investigation & formal analysis for Figure S1C&D.

**KO’H:** Funding acquisition, resources, supervision, methodology (building the light patterning system, setting up optogenetics pipeline)

**MOL:** Supervision, resources, methodology (building the light patterning system)

**CEB:** Conceptualisation, funding acquisition, resources, supervision, investigation & data curation (optogenetics imaging experiments), methodology (setting up optogenetics pipeline), writing – original draft, writing – review and editing.

## Figure Legends

**Movie 1**

Reversible deformation of the basal edge of the neural keel following RhoA activation

**Movie 2**

Velocity vector field of a 15 somite stage embryo undergoing RhoA activation demonstrates asymmetric anterior displacement

**Movie 3**

Velocity vector field of a 14 somite stage embryo undergoing RhoA activation demonstrates asymmetric posterior displacement

