## Supplementary Information for "Local optogenetic NMYII activation within the zebrafish neural rod results in long-range, asymmetric force propagation"

**Supplementary methods**

***ARRIVE experimental design***

Since the experimental units were either individual cells or regionally specific areas of tissue, internal controls were used; e.g. a non-optogenetic region of tissue or a region of tissue before optogenetic activation. Sample sizes are included in figure legends. Only embryos that were morphologically normal were included in the study. It is not possible to select for gender, so a random mixture of males and females were used. Blinding was not relevant to these experiments. However, embryos were selected in an unbiased way, based on their peak velocity (see methods). PIV data was then automatically generated from standardised regions of interest, as described in the figure legends. Outcome measures are described in the figures and legends. Statistics, details of transgenic animals and experimental procedures are detailed in the methods and below. Results are reported with variance and statistical outcomes, as detailed in figures and legends.

***Zebrafish Care***

Embryos were collected and staged according to standard protocols (Kimmel *et al*, 1995; Westerfield, 2000). Breeding stocks of adult zebrafish were bred and housed in a dedicated centralised zebrafish facility. Up to 14 adult fish were kept in 2.8L tanks. Fry were fed equal parts of Gemma Skretting 75, Skretting 150 and Freeze fried rotifers. Juvenilles were fed equal parts of Skretting 150, Skretting 300 and Golden pearl. Adults were fed (per 160g) 100g Skretting 500, 20g Golden pearl and 40g Spirulina flake. Animals over 14 d.p.f. were fed brine shrimp for alternate feedings. Water quality was carefully controlled to a PH of 7, conductivity of 600 and temperature of 27°C.

***Fusion constructs and mRNA microinjection***

Plasmids were constructed by Gibson assembly (Gibson *et al*, 2009). Genes were spaced using sequence encoding 10 amino-acid guanine-serine rich flexible linkers. DimericTomato-2xrGBD (Mahlandt *et al*, 2021) was inserted into a pCS2+ backbone for mRNA synthesis. mRNA was synthesised from pCS2+ plasmids linearised with NotI (Promega, R643A), using the SP6 mMessage mMachine kit (Ambion, AM1340). mRNA was diluted in nuclease-free water and injected into one cell of 4-32 cell embryos depending on the level of mosaicism desired (see Table 1 for mRNA amounts). For optogenetic experiments, 3-3.5 mM PCB (Sichem, SC-1800) dissolved in DMSO was injected with the mRNA, with these injections carried out in the dark.

*Table 1 - Fusion Constructs*

| **Name** | **Referred to in text** | **Reference** | **mRNA injected (ng)** |
|---|---|---|---|
| pCS2-N-PAS2-GAF-PHYB-mCherry-CAAX | PhyB-mCherry-Caax | Addgene plasmid #154910, (Buckley *et al*, 2016) | 0.312 - 0.585 |
| pCS2-N-PAS2-GAF-PHYB-CAAX | PhyB-caax | This study | 0.26 |

| | | | |
|---|---|---|---|
| pT3TS-Tol2 | Transposase RNA | pT3TS-Tol2 was a gift from Stephen Ekker (Addgene plasmid # 31831; http://n2t.net/addgene:31831 ; RRID:Addgene_31831) (Balciunas *et al*, 2006) | 0.025 |
| ubb:Ath.Pif6-EGFP-arhgef12a | Pif6-EGFP-LARG | This study | - |
| pCS2+-H2B-RFP | H2B-RFP | (Buckley *et al*, 2013) | 0.026 |
| CMVdel-dTomato-2xrGBD | dimericTomato-2xrGBD | dimericTomato-2xrGBD was a gift from Dorus Gadella (Addgene plasmid # 176098; http://n2t.net/addgene:176098 ; RRID:Addgene_176098) (Mahlandt *et al.*, 2021) | - |
| pCS2+-dTomato-2xrGBD | dT-2xrGBD | This study | 0.0364 - 0.0468 |

***Transgenic line generation***
2.5pg of the *ubb:Ath.Pif6-EGFP-arhgef12a* plasmid was injected into the cell of flat 1-cell stage, Tüpfel (TL) zebrafish embryos, along with 25pg of transposase RNA, generated from the pT3TS-Tol2 plasmid using the T3 mMessage mMachine kit (Ambion, AM1348). Embryos were screened for EGFP fluorescence and possible founders were grown to adulthood. These were then outcrossed to wild-type zebrafish. F1 embryos positive for EGFP fluorescence were grown to adulthood. These were then outcrossed to wild-type zebrafish and the proportion of EGFP embryos for each F1 fish was assessed. We selected an F1 fish that generated 50% fluorescent embryos (as expected for a single insertion of the transgene), had ubiquitous, medium-fluorescence expression within the hindbrain, and responded well to optogenetic activation. This was used to generate subsequent generations of the stable line.

***Confocal Unit***
The imaging part of the photomanipulation system is built around a Nikon ECLIPSE Ti2 inverted microscope with incubation system and a commercial disc confocal system (CrestOptics X-light V3). The confocal system is composed of an LDI-7 Laser Diode Illuminator (89 North), a lenslet array, a moveable spinning disk, two cameras (Photometrics Prime 95B), filters and lenses. The excitation light from the Laser illuminator is cleaned by an excitation filter (Chroma ZET405/470/555/640x). Then the light is focused on the spinning disk (CrestOptics, SP 50 µm/400 µm pinhole), passing through the pin holes, reflected by a dichroic mirror (Chroma ZT405/470/555/640rpc-UF2), and refocused on the microscope imaging plane. The emission light from the specimen is collected by the same objective, following the same beam path to the spinning disk, cleaned by an emission filter, (Chroma ZET405/470/555/640m-OD8), and projected to the camera. The emission light can be split into 2 channels with a dichroic mirror (Chroma T565lpxr-UF2). A matching bandpass filter is placed in front of each camera to limit the detected spectral range and block unwanted stray lights (Chroma ET525/50m and ET605/70m). The typical excitation power density at the sample plane for imaging were around 6 mW/cm$^2$ for the 470 nm laser (at 25%) and 0.4 mW/cm$^2$ for the 555 nm laser (at 1%).

A collimated 730 nm LED (Thorlabs M730L5) was mounted within the incubation system for optogenetic deactivation of the whole imaging dish between experiments. For all imaging, a 40x NA 1.25 Silicone objective (Nikon CFI Plan Apochromat Lambda S 40XC Sil) was used.

***Immunohistochemistry***
Embryos were fixed for 3 hours in 4% paraformaldehyde, washed in 0.1% Triton in PBS (PBST), and blocked for 1 hour with 10% goat serum at room temperature. Embryos were stained overnight with rabbit anti-Phospho-Myosin Light Chain 2 (Thr18/Ser19) (Cell Signalling Technology, 3674) at 1:50 dilution at 4°C. They were then washed in PBST and stained with goat anti-rabbit Atto 647N (Sigma-Aldrich 40839) at 1:500 dilution and Hoescht at 1:1000 dilution for 3 hours at room temperature, then washed again.

***Image analysis***
Image analysis was performed in Fiji, version 2.14.0, unless otherwise described. Line profiles were 4 pixels (1.1µm) wide and performed using the plot profile function. Membrane and cytoplasmic regions were selected using PhyB-mCherry-CAAX labelling or recruited PIF6-EGFP-LARG. Rhombomere anterior-posterior lengths were calculated from the mean of 3 measurements. Nuclear displacement was quantified by the difference between centroids of manually outlined nuclei in the north or south regions at the timepoint 60s before activation and immediately before activation, immediately before activation and 60s after activation, and immediately before deactivation and 60s after deactivation.

**Supplementary Figure 1**
**Rapid optogenetic LARG membrane recruitment results in RhoA activation and MLC2 phosphorylation in zebrafish neuroepithelial cells**

A. Representative images showing dimericTomato-2xrGBD RhoA biosensor (magenta) and PIF6-EGFP-LARG (cyan) recruitment upon 730nm and 640nm illumination, indicating activation of RhoA under 640nm illumination. Scale bar 10µm
B. Average line profiles of PIF6-EGFP-LARG (green) and dimericTomato-2xrGBD (magenta) across cell-cell interfaces upon 730nm (dotted) and 640nm (solid) illumination, normalised to the membrane intensity during 730nm illumination. Graph shows mean and SEM. N = 59 cell-cell interfaces from 5 embryos from 3 experiments
C. Representative sum intensity projections showing doubly phosphorylated MLC2 (ppMLC2 immunohistochemistry, magenta) upregulation and PIF6-EGFP-LARG (cyan) recruitment in a PhyB-CAAX positive ROI (marked by H2B-RFP positive nuclei (yellow)), and lack of ppMLC2 and PIF6-EGFP-LARG recruitment in a PhyB-CAAX negative adjacent ROI after 640nm illumination. Scale bar 10µm.
D. Mean intensity of ppMLC2 after 640nm illumination in PhyB-CAAX positive (yellow) and negative (grey) ROIs. Boxplots show median, points indicate ROIs, N = 7 PhyB-CAAX positive ROI and 7 PhyB-CAAX negative ROIs in 2 embryos from 1 experiment. Mean intensity was significantly different between groups using two-sample t-test ($t(12) = 2.34$, *$p = 0.0374$).
E. PIF6-EGFP-LARG membrane/cytoplasmic intensity over time following 640nm and 730nm illumination in the ROI. Ratios were normalised to levels at t = 0s of 640nm illumination. Data points are mean ± SEM of N = 10 embryos, curves are fits to non-linear models of exponential association and dissociation with the ribbon showing the 95% CI, $y = 1.018 + (1.588 - 1.018) * (1 - e^{(-0.083 * x)})$ and $y = 1.126 + (1.673 - 1.126) * e^{(-0.093 * x)}$ for 640nm and 740nm illumination respectively. The rate constants are -0.083 and -0.093 for 640nm and 740nm illumination respectively. Time constants (τ) are the inverse of the rate constant and indicate the time taken for the membrane/cytoplasmic intensity to increase by a factor of $1 - 1/e$ in the case of 640nm illumination, or decrease by a factor of $1/e$ in the case of 730nm illumination.

**Supplementary Figure 2**
**Simulated 3D light patterning using a 15µm x 15µm ROI**

A. Simulated cross section (xz) of the 640nm activation (i) and 730nm deactivation (ii) illumination intensity resulting from a 15µm x 15µm ROI. Red line indicates 70% intensity.
B. Intensity line profile of simulated activation/deactivation light at activation plane and at 10µm intervals away from the activation plane for a 15µm x 15µm ROI.

**Supplementary Figure 3**
**Lack of response in region opposite to direction of tissue movement.**

A.
  i. Average normalised velocity (relative to the preactivation mean) in the 'south' region of anteriorly directed embryos, n = 6.
  ii. Average normalised velocity (relative to the preactivation mean) in the 'north' region of posteriorly directed embryos, n = 3.

B.
  i. Average velocity direction (relative to the preactivation mean) in the 'south' region of anteriorly directed embryos, n = 6
  ii. Average velocity direction (relative to the preactivation mean) in the 'north' region of posteriorly directed embryos, n = 3

**Supplementary Figure 4**
**Oscillations in average isotropic strain rate before, during and after activation**.

A. Isotropic strain rate z-score in the anterior-posterior column before RhoA activation.
B. Isotropic strain rate z-score in the anterior-posterior column during RhoA activation in the ROI.
C. Isotropic strain rate z-score in the anterior-posterior column after RhoA activation.
   Average of anteriorly directed embryos, n = 6. Each line indicates a timepoint.

D. Quantification of anterior-posterior length of rhombomeres 4 and 5 in neural rod stage embryos (n = 12 embryos, 17-19 somite stage). Average length of R4 is 60.6 ± 5.3 µm and R5 is 60.1 ± 4.7 µm (mean ± s.d.). Boxplots show median, points indicate individual embryos.

**Supplementary Figure 5**
**Peak strain rates in posteriorly directed embryos**

A. Average isotropic strain rate z-score in the anterior-posterior column at the time of peak velocity during activation (average of posteriorly directed embryos, n = 3).
B. Average anisotropic strain rate z-score in the anterior-posterior column at the time of peak velocity during activation (average of posteriorly directed embryos, n = 3).
C. Raw anisotropic (pink) and isotropic (blue) strain rates in the anterior-posterior column at the time of peak velocity during activation (average of posteriorly directed embryos, n = 3).

**Supplementary Figure 6**
**Gradients in Pif6-EGFP-LARG recruitment may be introduced by preferential recruitment to the tissue putative midline**

A. Representative diagram showing membrane regions measured in the ROI during activation for Pif6-EGFP-LARG expression and before the timelapse experiment for PhyB-mCherry-Caax expression for B and C.
B. Gradients of Pif6-EGFP-LARG membrane/cytoplasm ratio across the AP axis of the ROI for anteriorly and posteriorly directed embryos. Linear regression lines in colour show individual embryos, slope indicates the mean slope for each group (n = 11 and n = 3 for anterior and posterior embryos respectively).

C. Lack of correlation between Pif6-EGFP-LARG gradient and PhyB-mCherry-Caax gradient. Points indicate individual embryos, colour matches with embryos in B. Spearman's rank correlation coefficient was not significant (rho = 0.086, p = 0.919).
D. Representative images of global optogenetic RhoA activation show preferential recruitment to cells undergoing mitosis (asterisk) and to the midline region (arrow head) in the neural keel and neural rod. High LARG recruitment can occur despite weak PhyB-mCherry-Caax expression.
E. RhoA biosensor expression is elevated at the midline of the neural tissue in the absence of optogenetic activation.
   i. Mean ± SEM of the intensity of RhoA biosensor taken across the mediolateral (ML) width of the neural tissue, n = 4 embryos from neural keel (n = 2) to neural rod (n = 2) stage.
   ii. Representative images of neural keel and neural rod RhoA biosensor expression – dotted regions indicate line profiles taken for i.

**Supplementary Figure 7**
**Experimental pipelines for optogenetic experiments**

A. Schematic of optogenetic timelapse experiment showing how imaging and digital mirror device (DMD) light patterning were combined to activate an ROI
B. Schematic of 3D activation experiment showing how a stack was built to show the changing levels of recruitment in the z-plane from a light-patterned ROI at a single plane

**Supplementary Movie 1**
Specific subcellular LARG recruitment to neuroepithelial progenitor cells with simultaneous 555nm (left) and 470nm (right) imaging of mosaic PhyB-MCh-Caax (left) and transgenic Pif6-Larg-EGFP (right)
