## Supplementary figures and images for "Local optogenetic NMYII activation within the zebrafish neural rod results in long-range, asymmetric force propagation"

### Figure S1

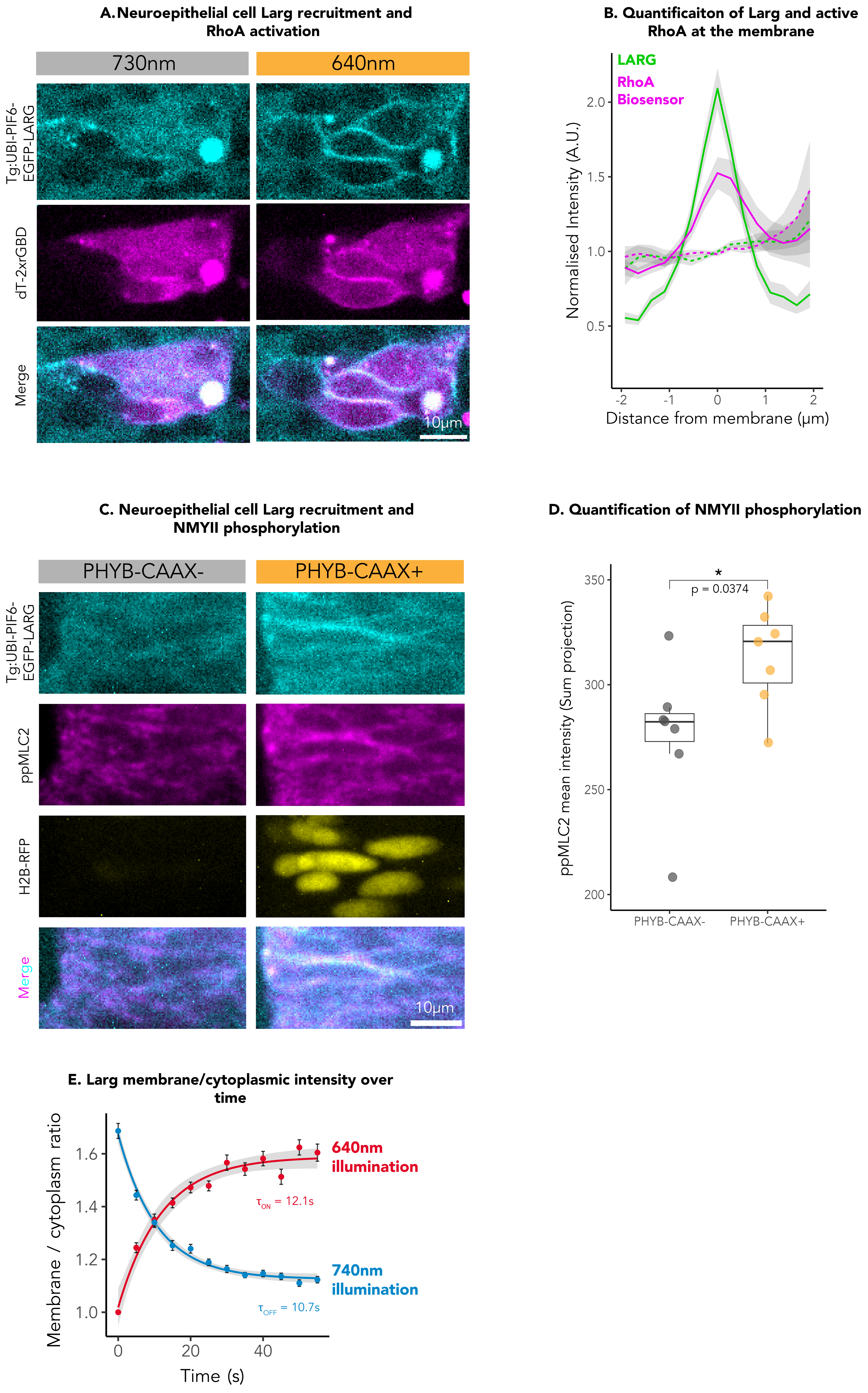

### Figure S3

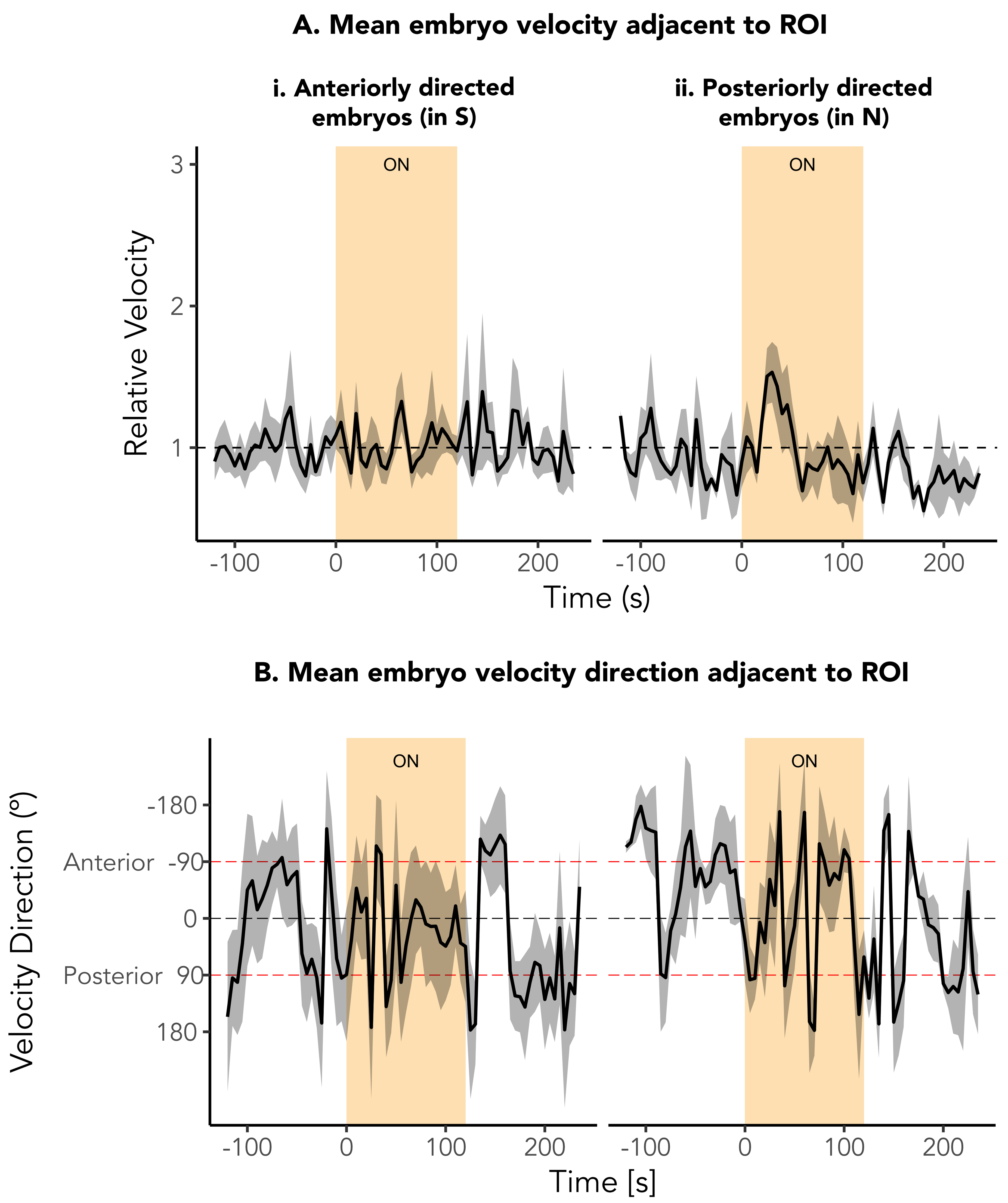

### Figure S6

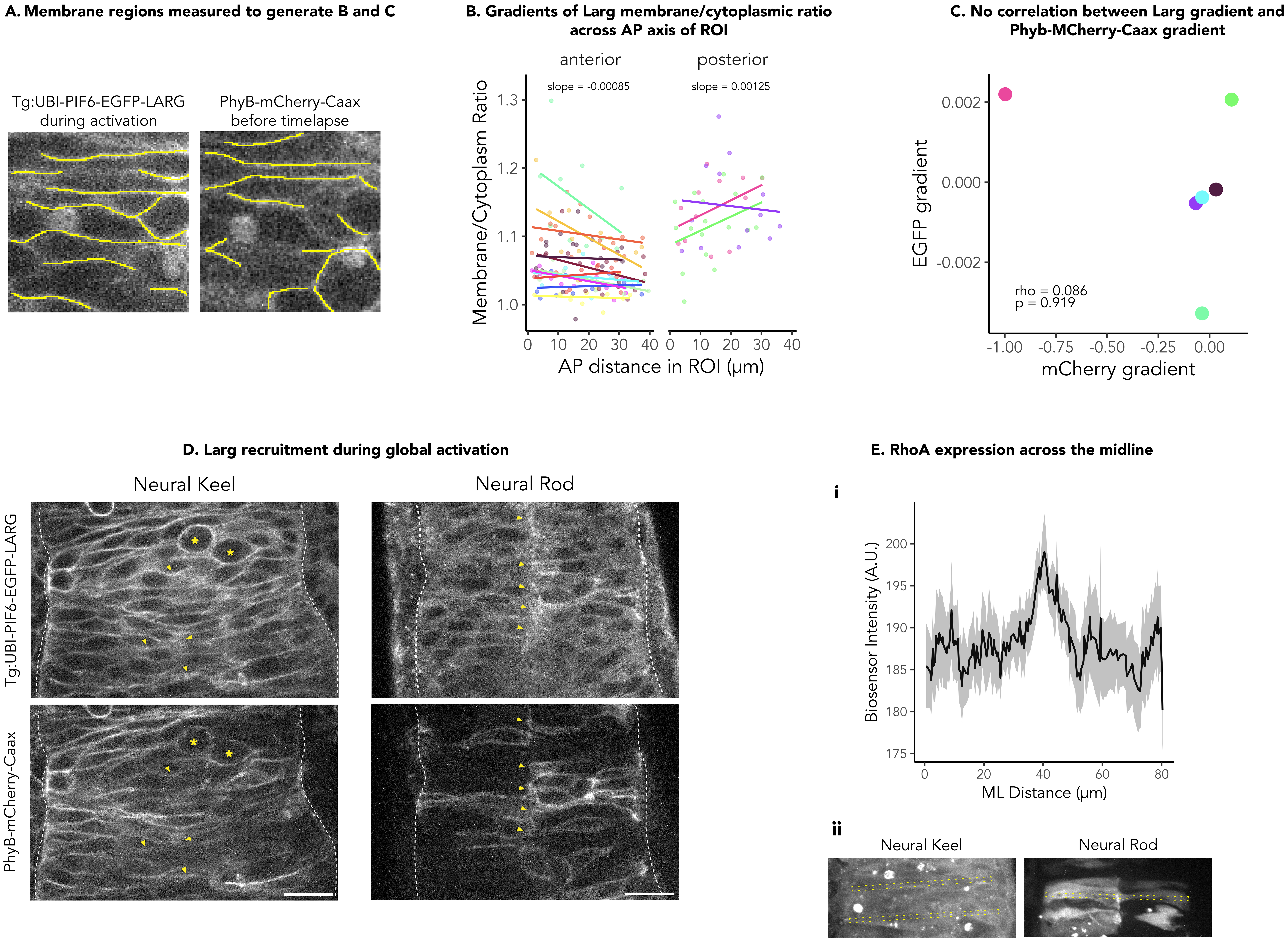

### Figure S7

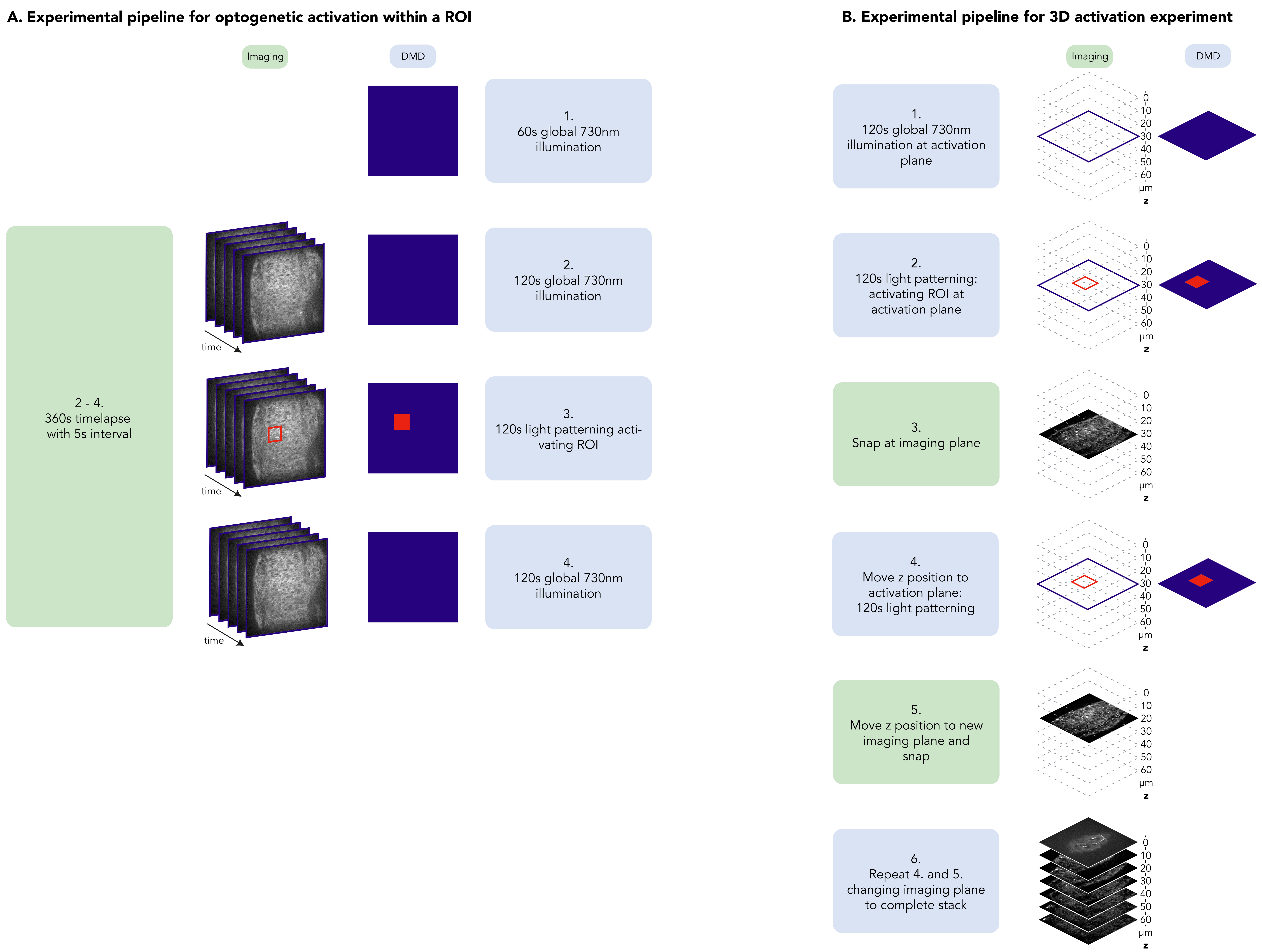
