## Supplementary material for "Local optogenetic NMYII activation within the zebrafish neural rod results in long-range, asymmetric force propagation": Figure S2

**A. Simulated cross-section of 640nm and 730nm light for a 15um x 15um ROI**

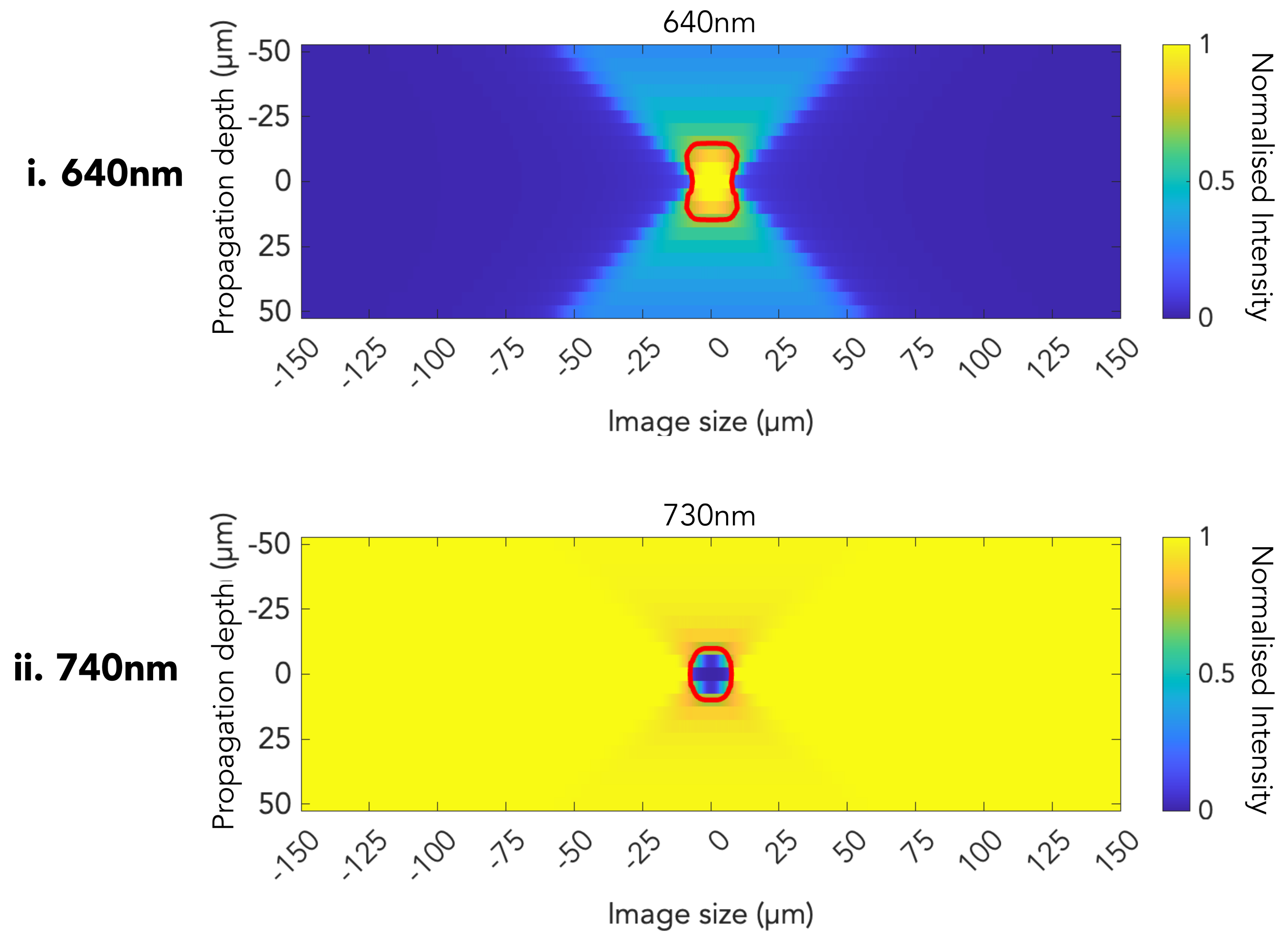


**B. Simulated intensity profile of 640nm and 730nm light at 10um intervals from ROI**

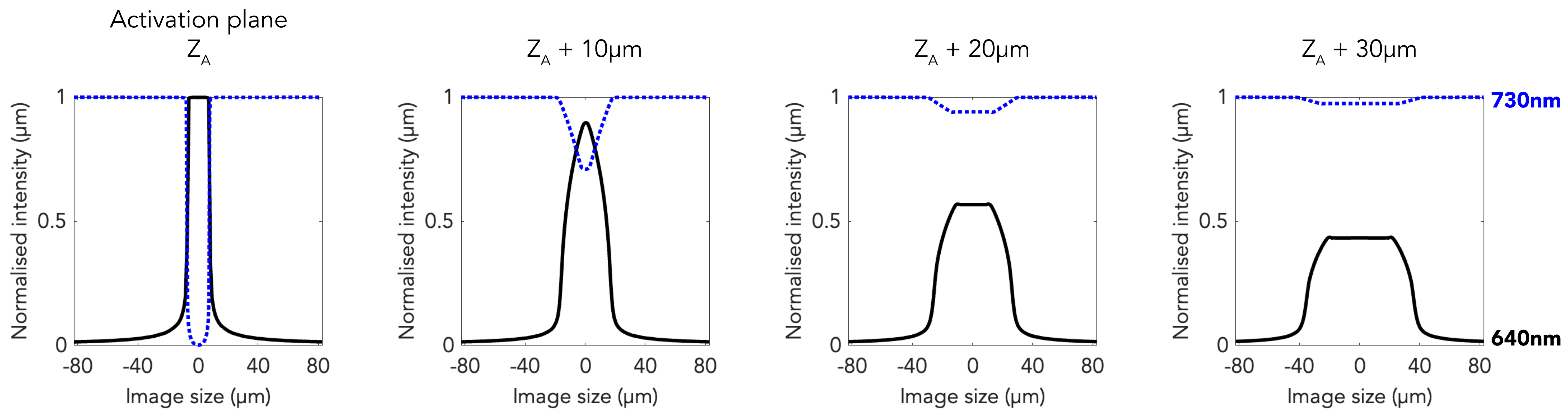
