## Supplementary material for "Local optogenetic NMYII activation within the zebrafish neural rod results in long-range, asymmetric force propagation": Figure S4

**Oscillations in average isotropic strain rate**

**A. Before RhoA activation**

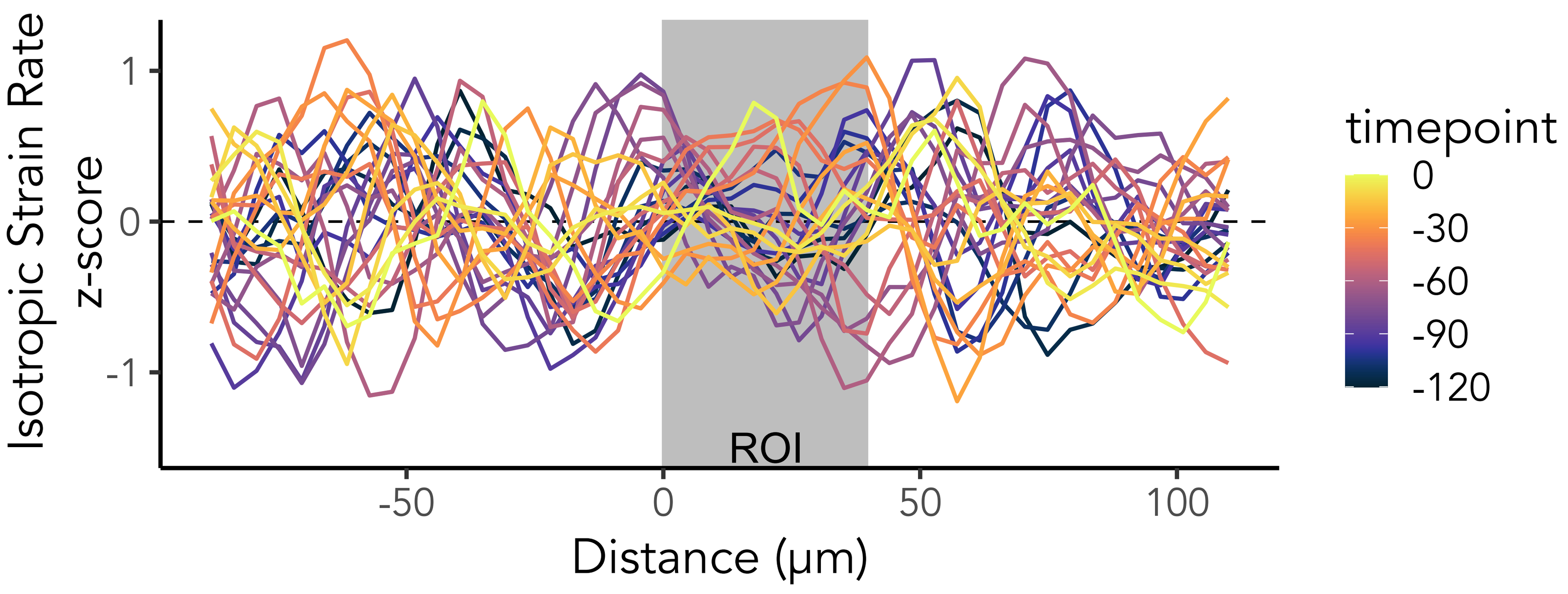


**B. During RhoA activation**

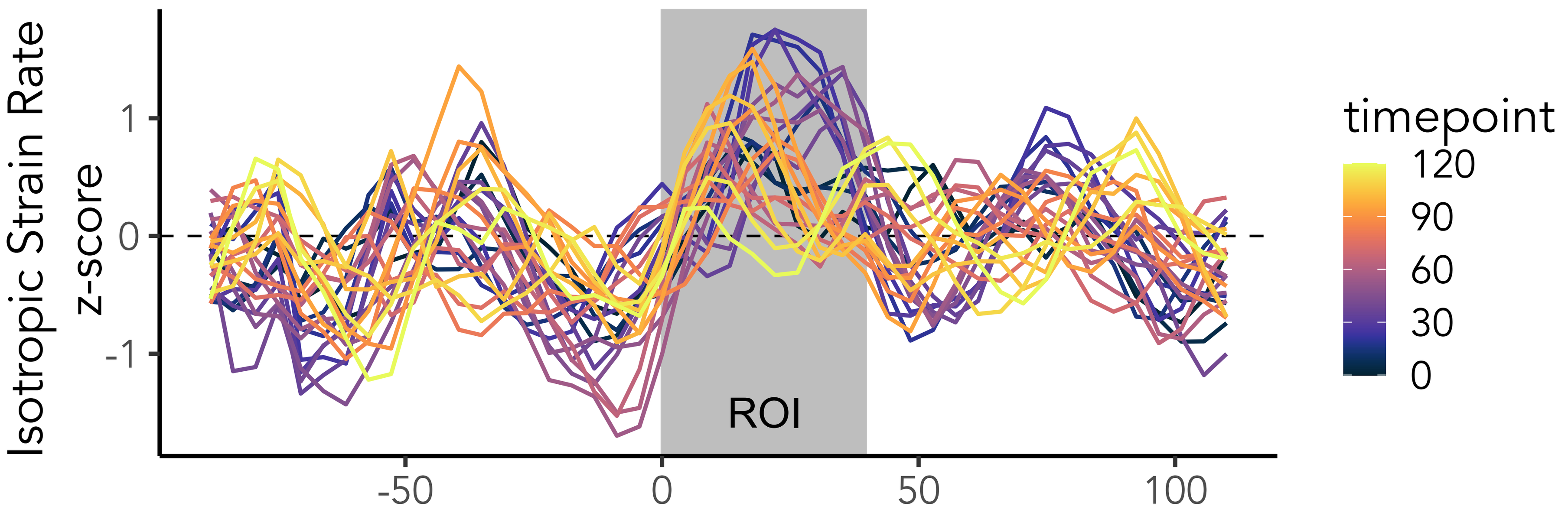


**C. After RhoA activation**

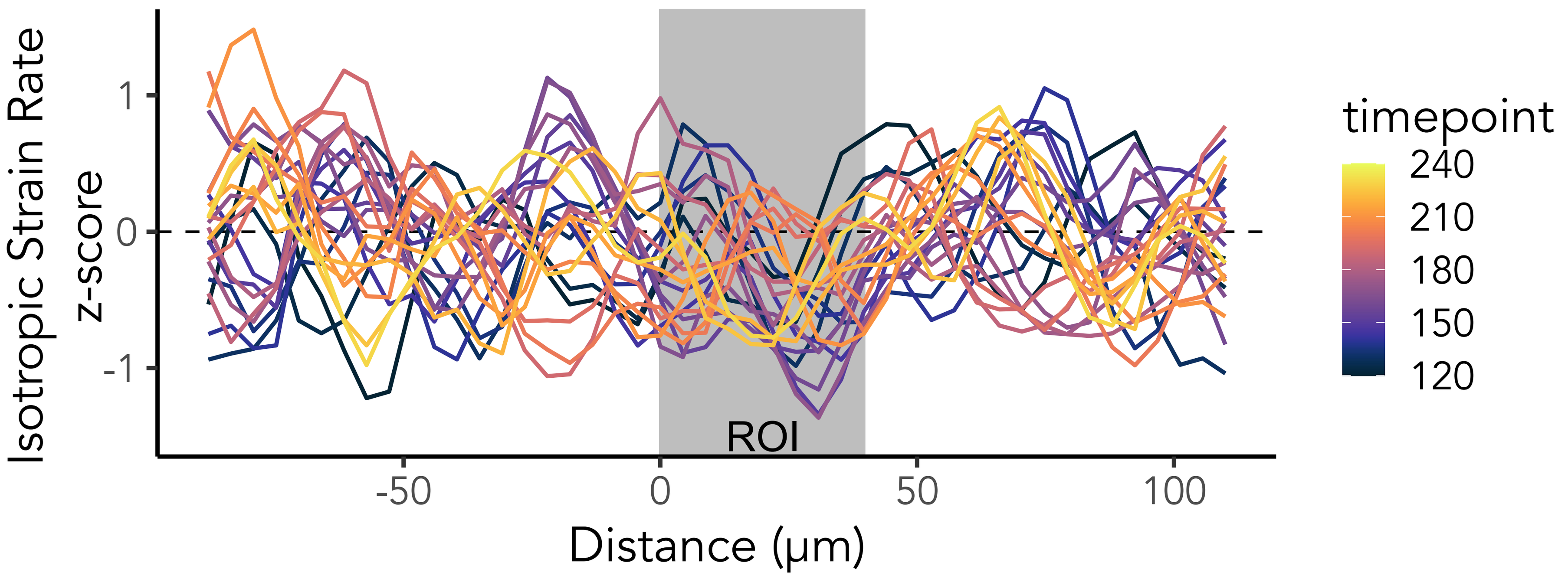


**Oscillation wavelength is within the range of rhombomere length**

**D. Rhombomere 4 and 5 anterior-posterior length**

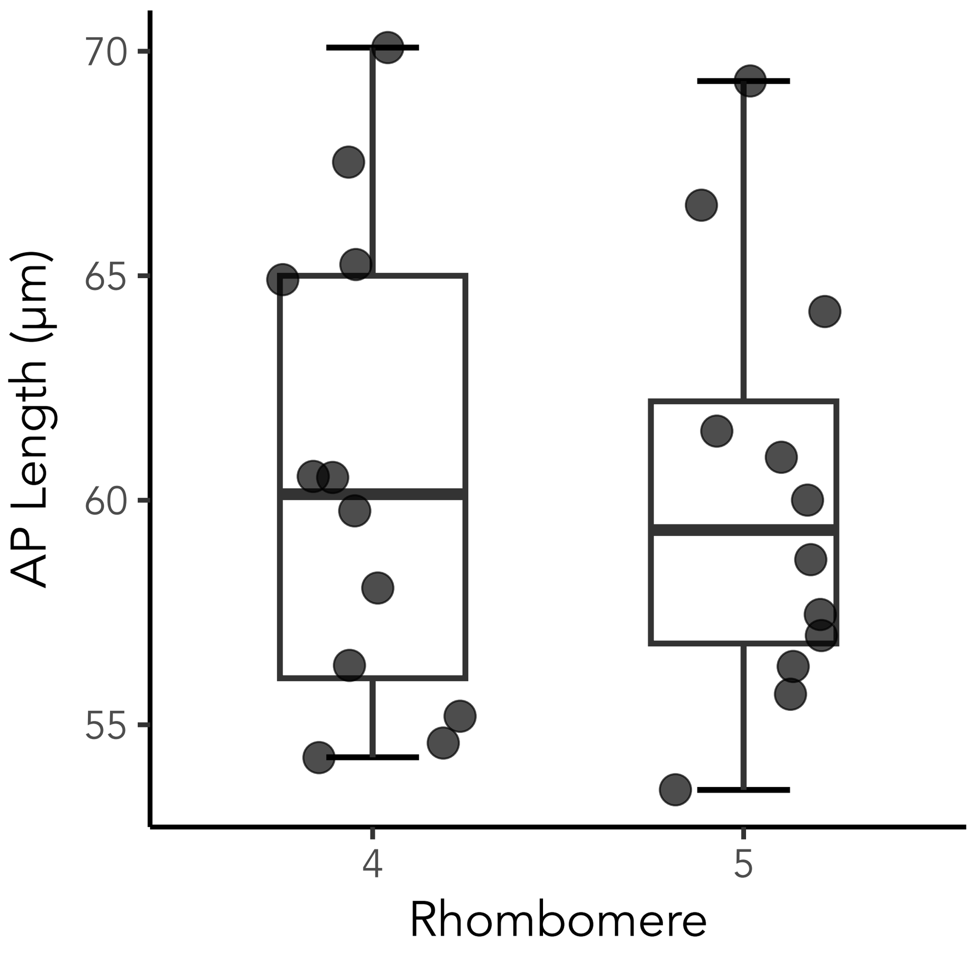
