## Supplementary material for "Local optogenetic NMYII activation within the zebrafish neural rod results in long-range, asymmetric force propagation": Figure S5

**A** **Distribution of mean isotropic strain rate during activation peak**

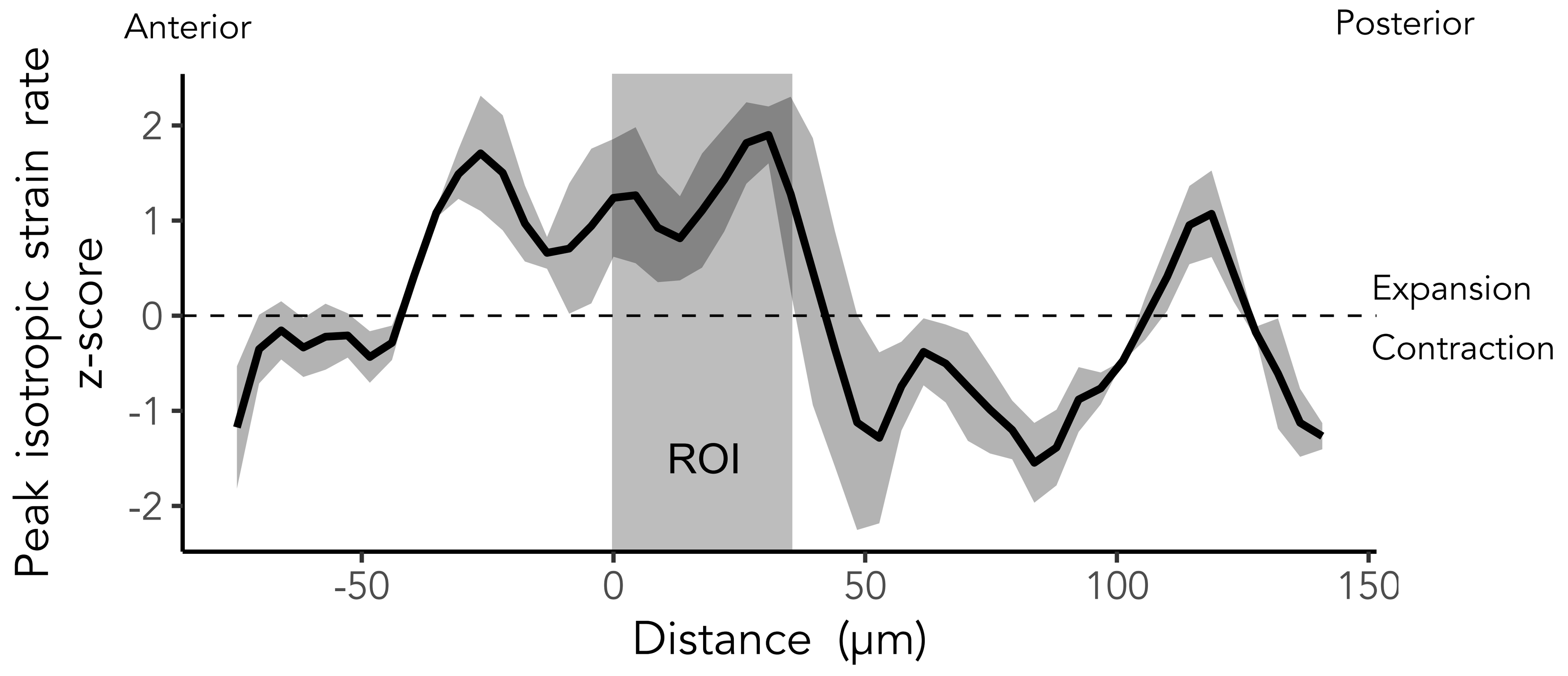


**B** **Distribution of mean anisotropic strain rate during activation peak**

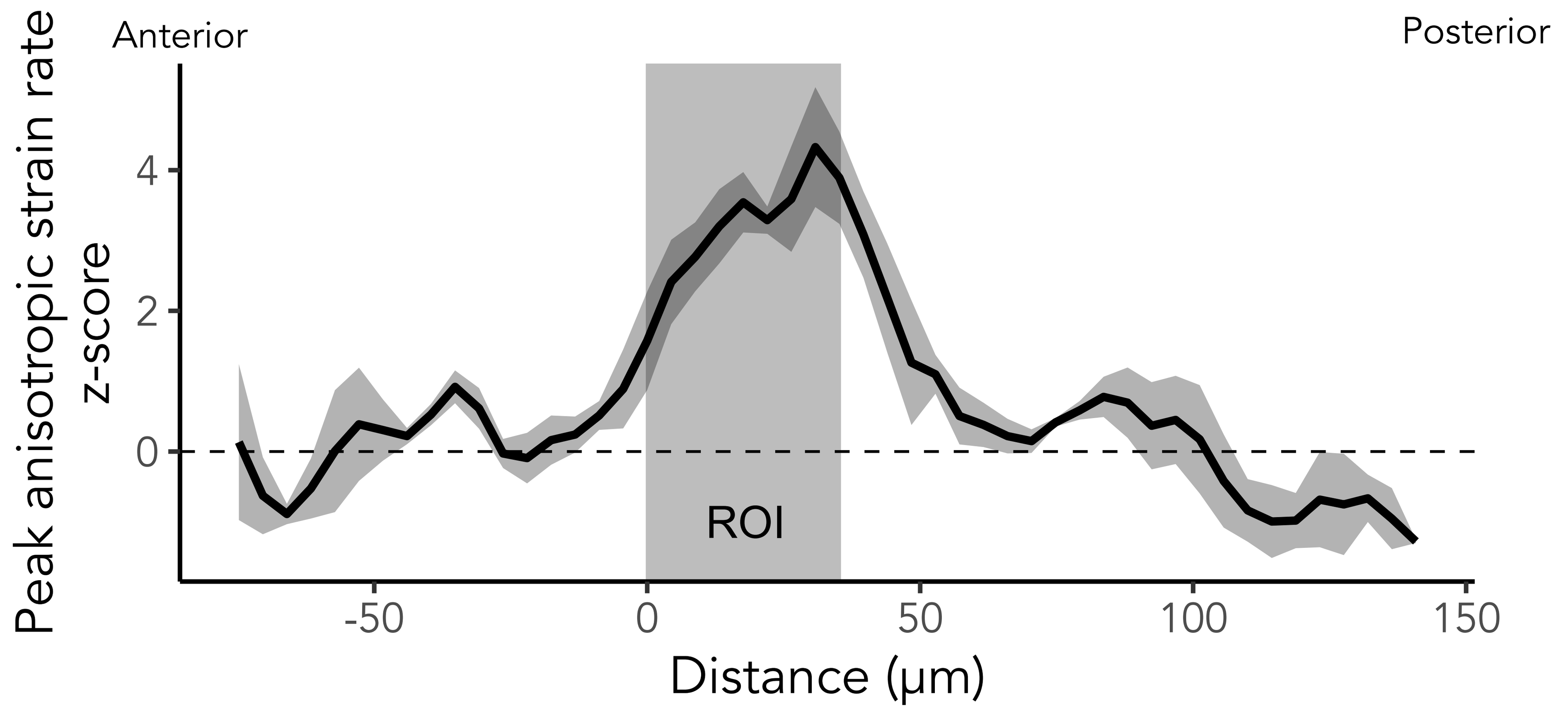


**C** **Comparison of mean strain rates**

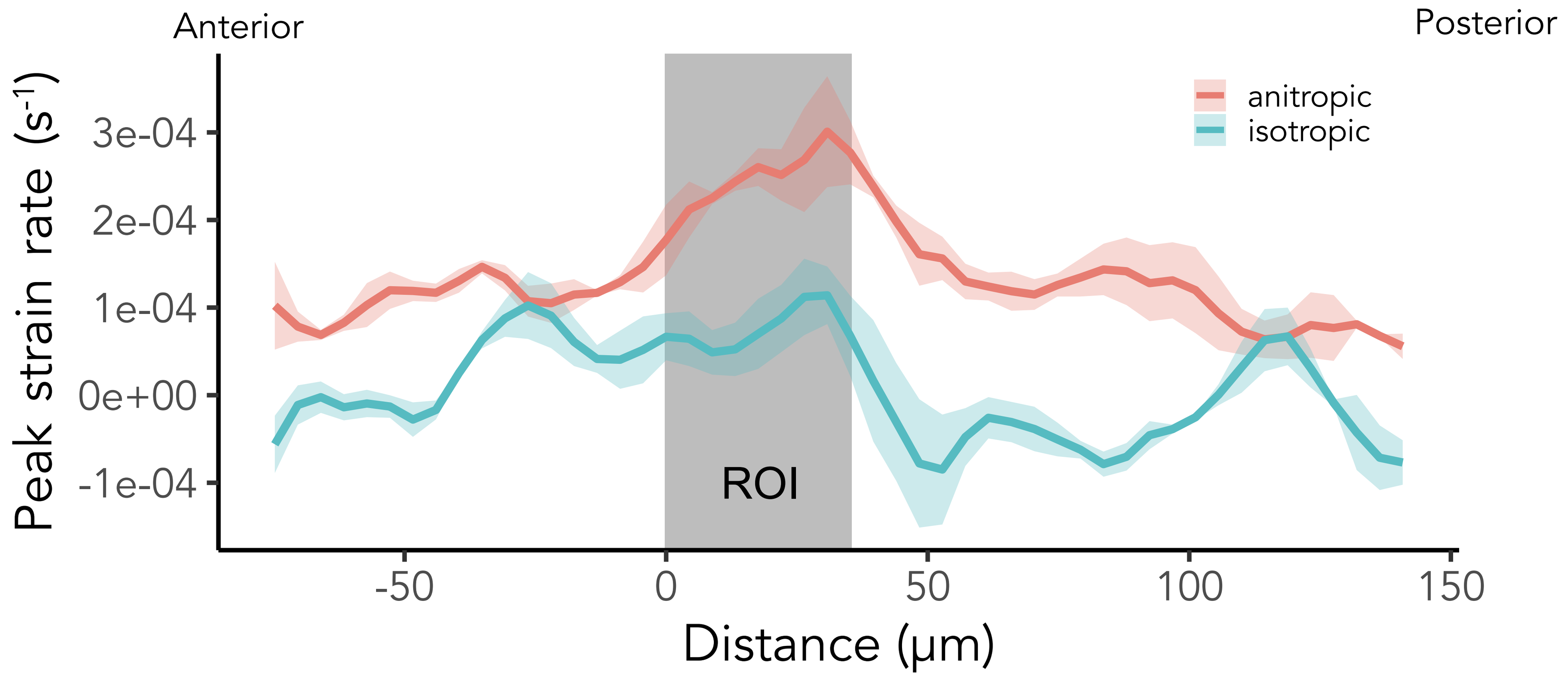
